# Functional characterization of the disease-associated *CCL2* rs1024611G-rs13900T haplotype: The role of the RNA-binding protein HuR

**DOI:** 10.1101/2023.10.31.564937

**Authors:** Feroz Akhtar, Joselin Hernandez Ruiz, Ya-Guang Liu, Roy G. Resendez, Denis Feliers, Liza D. Morales, Alvaro Diaz-Badillo, Donna M. Lehman, Rector Arya, Juan Carlos Lopez-Alvarenga, John Blangero, Ravindranath Duggirala, Srinivas Mummidi

**Affiliations:** Department of Health and Behavioral Sciences, Texas A&M University-San Antonio, Texas, USA; Utah Center for Genetic Discovery, Department of Human Genetics, University of Utah, Salt Lake City, Utah, USA; Department of Pathology, School of Medicine, University of Texas Health San Antonio, San Antonio, Texas, USA; Department of Medicine, School of Medicine, University of Texas Health San Antonio, San Antonio, Texas, USA; South Texas Diabetes and Obesity Institute, Department of Genetics, School of Medicine, University of Texas Rio Grande Valley, Brownsville, Texas, USA; Department of Population Health and Biostatistics, School of Medicine, University of Texas Rio Grande Valley, Harlingen, Texas, USA

## Abstract

CC-chemokine ligand 2 (CCL2) is involved in the pathogenesis of several diseases associated with monocyte/macrophage recruitment, such as HIV-associated neurocognitive disorder (HAND), tuberculosis, and atherosclerosis. The rs1024611 (alleles: A>G; G is the risk allele) polymorphism in the *CCL2 cis*-regulatory region is associated with increased CCL2 expression in vitro and ex vivo, leukocyte mobilization in vivo, and deleterious disease outcomes. However, the molecular basis for the rs1024611-associated differential CCL2 expression remains poorly characterized. It is conceivable that genetic variant(s) in linkage disequilibrium (LD) with rs1024611 could mediate such effects. Previously, we used rs13900 (alleles: C>T) in the *CCL2* 3’ untranslated region (3’ UTR) that is in perfect LD with rs1024611 to demonstrate allelic expression imbalance (AEI) of *CCL2* in heterozygous individuals. Here we tested the hypothesis that the rs13900 could modulate *CCL2* expression by altering mRNA turnover and/or translatability. The rs13900 T allele conferred greater stability to the *CCL2* transcript when compared to the rs13900 C allele. The rs13900 T allele also had increased binding to Human Antigen R (HuR), an RNA-binding protein, in vitro and ex vivo. The rs13900 alleles imparted differential activity to reporter vectors and influenced the translatability of the reporter transcript. We further demonstrated the role of HuR in mediating allele-specific effects on CCL2 expression in overexpression and silencing studies. Our studies suggest that the differential interactions of HuR with rs13900 could modulate CCL2 expression and could in part explain the interindividual differences in CCL2-mediated disease susceptibility.

## Introduction

Identification of functional and/or causal genetic variants continues to pose a significant challenge in the post-genome-wide association studies (post-GWAS) era (MacArthur et al., 2017; Tam et al., 2019; Visscher et al., 2017). While emphasis has been placed on polymorphisms that map to *cis*-elements in enhancers and promoters, genetic variants that disrupt cis-elements in RNA binding protein (RBP) motifs in 3’ UTR have received much less attention. For many genes, the interactions of their 3’ UTRs with specific stabilizing and destabilizing RBPs play a critical role in modulating post-transcriptional events such as mRNA turnover and translatability (Mayr, 2019; Schwerk and Savan, 2015). The lack of a mechanistic understanding of how SNPs localizing to RBP motifs could impact gene expression impedes a greater understanding of the variability in disease susceptibility and outcomes.

Post-transcriptional mechanisms are thought to play a crucial role in the initiation and resolution of the inflammatory response (Anderson, 2010; Yoshinaga and Takeuchi, 2019). Such regulation can profoundly affect gene expression levels; modest changes in mRNA stability can lead to significant effects on mRNA and protein abundance (Buccitelli and Selbach, 2020; Ross, 1995). Notably, a genome-scale study using mouse dendritic cells demonstrated that post-transcriptional mRNA degradation was a salient feature of inflammatory and immune signaling genes, as well as targets of NF-kappaB signaling following lipopolysaccharide (LPS) stimulation (Rabani et al., 2011). In addition, polymorphisms in non-coding regions such as the 3’ UTR can have a significant impact on mRNA stability and translatability. For example, a genome-wide study of variation in gene-specific mRNA decays in lymphoblastoid cell lines across individuals found about 195 genetic variants that are specifically associated with variation in mRNA decay rates, called “rdQTLs” (RNA Decay Quantitative Trait Loci) (Pai et al., 2012). The authors estimated that 35% of the most significant eQTL (Expression Quantitative Trait Loci) single nucleotide polymorphisms (SNPs) are associated with decay rates (Pai et al., 2012). In another study, Duan *et al*. reported that ∼37% of gene expression differences among individuals may be attributed to RNA half-life differences (Duan et al., 2013). Farh et al., reported that the 3’ UTRs are highly enriched for eQTL candidate causal SNPs (>1,500) relative to other transcribed SNPs (Farh et al., 2015). However, the molecular mechanisms by which these polymorphisms alter mRNA stability or translatability are poorly understood.

CCL2 is a potent monocyte chemoattractant produced by various cell types, either constitutively or following activation. CCL2 expression can be regulated by inflammatory molecules (e.g., IL-1, TNFα, LPS, and IFNγ) and growth factors (e.g., PDGF). While the cells of monocyte-macrophage lineage are a major source of CCL2 (Yoshimura et al., 1989), other cell types, such as fibroblasts, astrocytes, epithelial, and endothelial cells, also are an important source (Deshmane et al., 2009). CCL2 mediates recruitment of monocytes, memory T-cells, and dendritic cells to the site of inflammation (Gschwandtner, Derler and Midwood, 2019; Melgarejo et al., 2009) and there is substantial evidence implicating CCL2 as a key mediator in macrophage-mediated diseases. Rovin et al. described a SNP in the 5’-regulatory region of *CCL2* annotated as rs1024611 (dbSNP database; originally designated as – 2518A>G (Rovin, Lu and Saxena, 1999) or –2578A>G (Gonzalez et al., 2002)) that was associated with increased plasma CCL2 expression (Rovin, Lu and Saxena, 1999). This SNP is associated with increased serum CCL2 levels, enhanced macrophage recruitment to tissues, and progression to HIV-associated dementia (Gonzalez et al., 2002). Other studies showed that the rs1024611 G allele is associated with increased CCL2 levels in the plasma, urine, and cerebrospinal fluid in health and disease, and in tissues such as liver and skin (Cho et al., 2004; Fenoglio et al., 2004; Joven et al., 2006; Letendre et al., 2004; McDermott et al., 2005). The rs1024611 polymorphism has been associated with several diseases, including myocardial infarction (McDermott et al., 2005), carotid atherosclerosis (Alonso-Villaverde et al., 2004), pulmonary tuberculosis (Flores-Villanueva et al., 2005), severe acute pancreatitis (Cavestro et al., 2010), lupus nephritis (Tucci et al., 2004), asthma susceptibility and severity (Szalai et al., 2001), Crohn’s disease (CD) (Palmieri et al., 2010), Alzheimer’s disease (Fenoglio et al., 2004), and infections by Japanese Encephalitis virus (Chowdhury and Khan, 2017) and SARS-CoV-1 (Tu et al., 2015). Notably, rs3091315, a GWAS risk variant for CD (Franke et al., 2010) and inflammatory bowel disease (IBD) (Liu et al., 2015) is in a strong linkage disequilibrium (LD) with both rs1024611 (D′=1.0, *r^2^*=0.98) and rs13900 (D′=1.0, *r^2^*=0.98) in the CEU population. Given the importance of disease associations with rs1024611, significant efforts have been made by other groups and us to understand the molecular basis of this differential CCL2 expression associated with this polymorphism (Gonzalez et al., 2002; Mummidi, Bonello and Ahuja, 2009; Page et al., 2011; Pham et al., 2012; Wright et al., 2008). However, these studies did not provide a mechanistic link between the rs1024611 polymorphism and CCL2 expression, giving rise to the possibility that SNPs in strong LD with rs1024611 could be mediating these effects.

To identify potential functional SNPs that could explain the variability in CCL2 expression, we developed an extensive LD map of the *CCL2* genomic locus and reported that a SNP designated as the rs13900 (NM_002982.4:c.*65=) in the *CCL2* 3’ UTR is in perfect LD with rs1024611 and can serve as its proxy (Pham et al., 2012). We and others have demonstrated AEI in *CCL2* using rs13900 as a marker with the T allele showing a higher expression level relative to C allele (Johnson et al., 2008; Pham et al., 2012). In this study, we show that the differential binding of Human Antigen R (HuR), an RBP previously implicated in CCL2 expression, leads to altered stability and translatability of *CCL2* transcripts providing a mechanistic explanation for increased CCL2 expression in individuals with the rs1024611G-rs13900T haplotype and inter-individual differences in disease susceptibility associated with this haplotype.

## Results

### Individuals heterozygous for rs13900 show AEI of *CCL2*

We and others have previously reported a perfect linkage disequilibrium between rs1024611 in the *CCL2* cis-regulatory region and rs13900 in its 3′ UTR and that rs13900 can serve as a proxy for the disease-associated rs1024611 (Hubal et al., 2010; Intemann et al., 2011; Kasztelewicz et al., 2017; Pham et al., 2012). We further showed that *CCL2* exhibits allelic expression imbalance (AEI) in heterozygous individuals, with the rs13900 T allele (alternative allele) having a higher expression than the rs13900 C allele (reference allele) (Figure 1A). For this study, we recruited 47 healthy unrelated individuals (18–35-year-old) who were screened for rs13900 (Figure S1). Supplementary Table 1 shows the genotype and allele frequencies for rs13900 polymorphism in these recruited individuals. We found that the rs13900 C allele was at a higher frequency than the rs13900 T allele and the genotype frequencies were in line with Hardy-Weinberg equilibrium (*P* ≥ 0.05).. We reconfirmed that *CCL2* exhibits AEI using data from heterozygous individuals. For this we used total RNA obtained from LPS treated PBMC as described previously (Pham et al., 2012). LPS induced *CCL2* expression in PBMCs as confirmed by qRT-PCR, with about a 4.3-fold increase at 1 h, 6.09-fold increase at 3 h, and 1.94-fold increase at 6 h (Figure S2). We used the 3 hour stimulation in the subsequent experiments as the peak *CCL2* expression was detected at this time point following LPS stimulation. AEI was measured by quantifying the relative amount of the two alleles i.e., alternative allele (T) to reference allele (C), measured from the chromatogram after normalization of peak intensity using PeakPeaker v.2.0 (Figure 1B and 1C). This strategy ensures a direct comparison between the amount of *CCL2* mRNA that is transcribed from each allele or haplotype and that each allele is equally subjected to the effects of any external factors. gDNA was utilized as a control. The boxplot illustrates a notable difference in the detected levels of the C and T allele in the gDNA and cDNA with a higher expression of T allele relative to C allele (*P* < 0.005) (Figure 1D).

**Figure 1.**
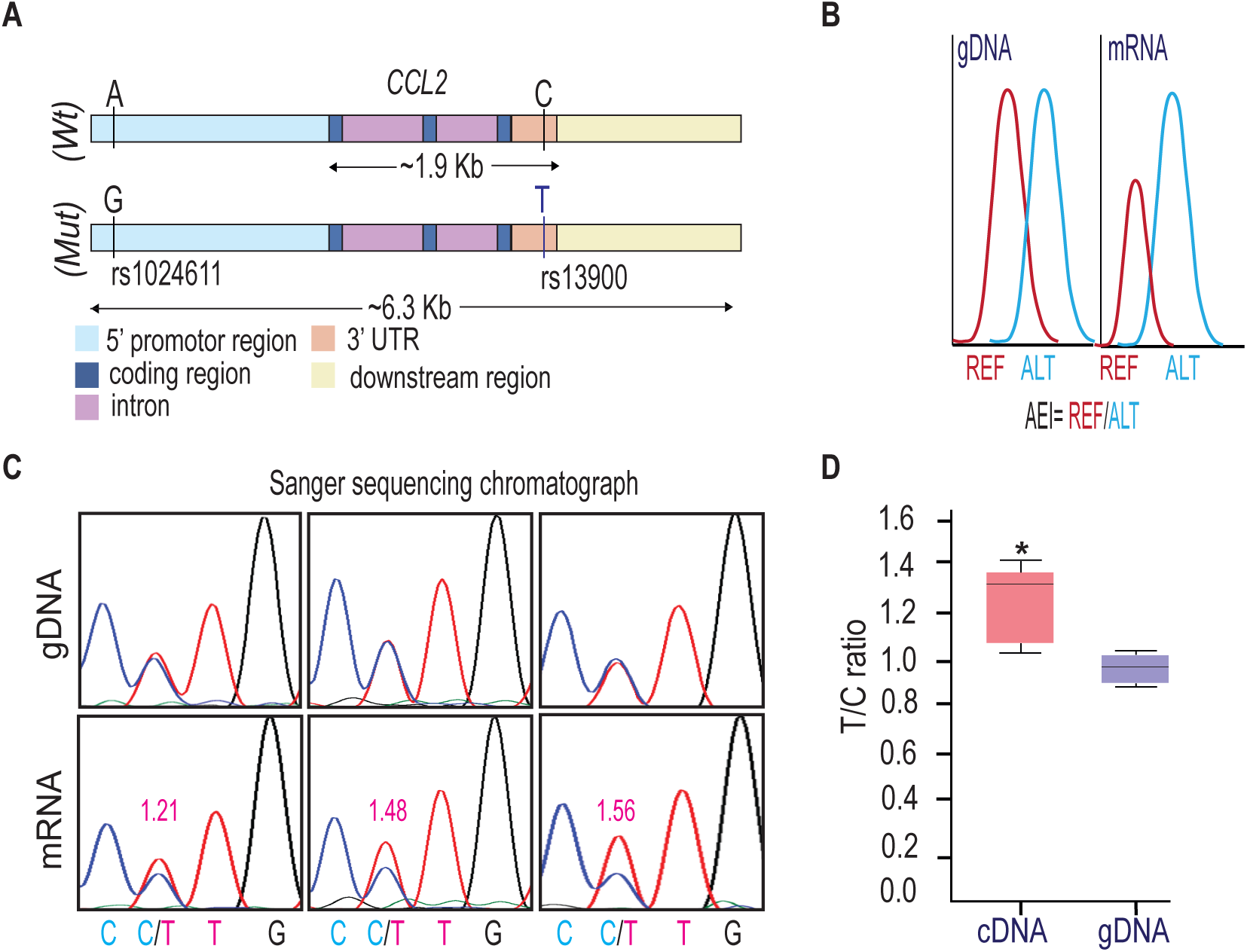
rs13900 heterozygous individuals exhibit AEI in *CCL2*. **(A)** Schematic depicting distal, proximal regulatory elements extending 3 kb on either side of *CCL2* gene and the LD between regulatory polymorphism rs1024611 and the transcribed polymorphism rs13900. rs1024611 is located 2578 base pairs upstream of the *CCL2* translation start site and rs13900 located in the *CCL2* 3′ UTR. **(B)** Allelic expression imbalance (AEI) in heterozygous donors is measured as a ratio of alternative allele (ALT) to reference allele (REF) in a transcribed polymorphism. **(C)** Representative chromatograms obtained following Sanger sequencing of PCR products obtained from genomic DNA (gDNA) and reverse transcription-PCR of mRNA (cDNA) from three individuals heterozygous for rs13900. gDNA and mRNA were obtained from PBMC treated with LPS for 3 h as previously described. The allelic ratios shown were determined by PeakPicker analysis. Peakpicker calculates allelic ratios by dividing the peak height of the alternate allele (rs13900 T allele) by that of reference allele (rs13900 C allele). The gDNA peaks were used for normalization. **(D)** Allelic ratio for cDNA and gDNA in six individuals heterozygous for rs13900 after treatment with LPS for 3 h. Statistical significance for the difference in the level of expression between the alleles was determined using Student’s *t*-test (*P* < 0.003).

### *CCL2* mRNA transcripts bearing rs13900 C and T alleles have different stability

Previous studies have shown that *CCL2* mRNA is subjected to post-transcriptional regulation through modulation of mRNA stability (Hao and Baltimore, 2009). Here, we determined whether the rs13900 modulates mRNA stability which may in part explain AEI. We used purified monocytes from four heterozygous individuals that were left untreated or stimulated with LPS for 3 h. Cells were either harvested after 3 h of LPS stimulation (considered as t=0) or cultured in the presence or absence of the transcriptional inhibitor actinomycin D for an additional 1, 2, or 4 h. Total RNA was isolated at each time point to assess *CCL2* transcript, calculated as fold induction over unstimulated cells. *CCL2* mRNA levels showed a strong upregulation following LPS stimulation (*P* < 0.05) (Figure 2A). Actinomycin D treatment revealed the kinetics of *CCL2* transcript degradation by determining the mRNA half-life, which represents the time (expressed in hours) at which mRNA expression is 50% of the initial level (Figure 2B; t_1/2_=Ln (0.5)/slope). For *CCL2*, t_1/2_=1.763 h and is in line with previously published studies that *CCL2* mRNA stability is modulated following inflammatory and cytokine stimuli (Hao and Baltimore, 2009; Zhai et al., 2008).

**Figure 2.**
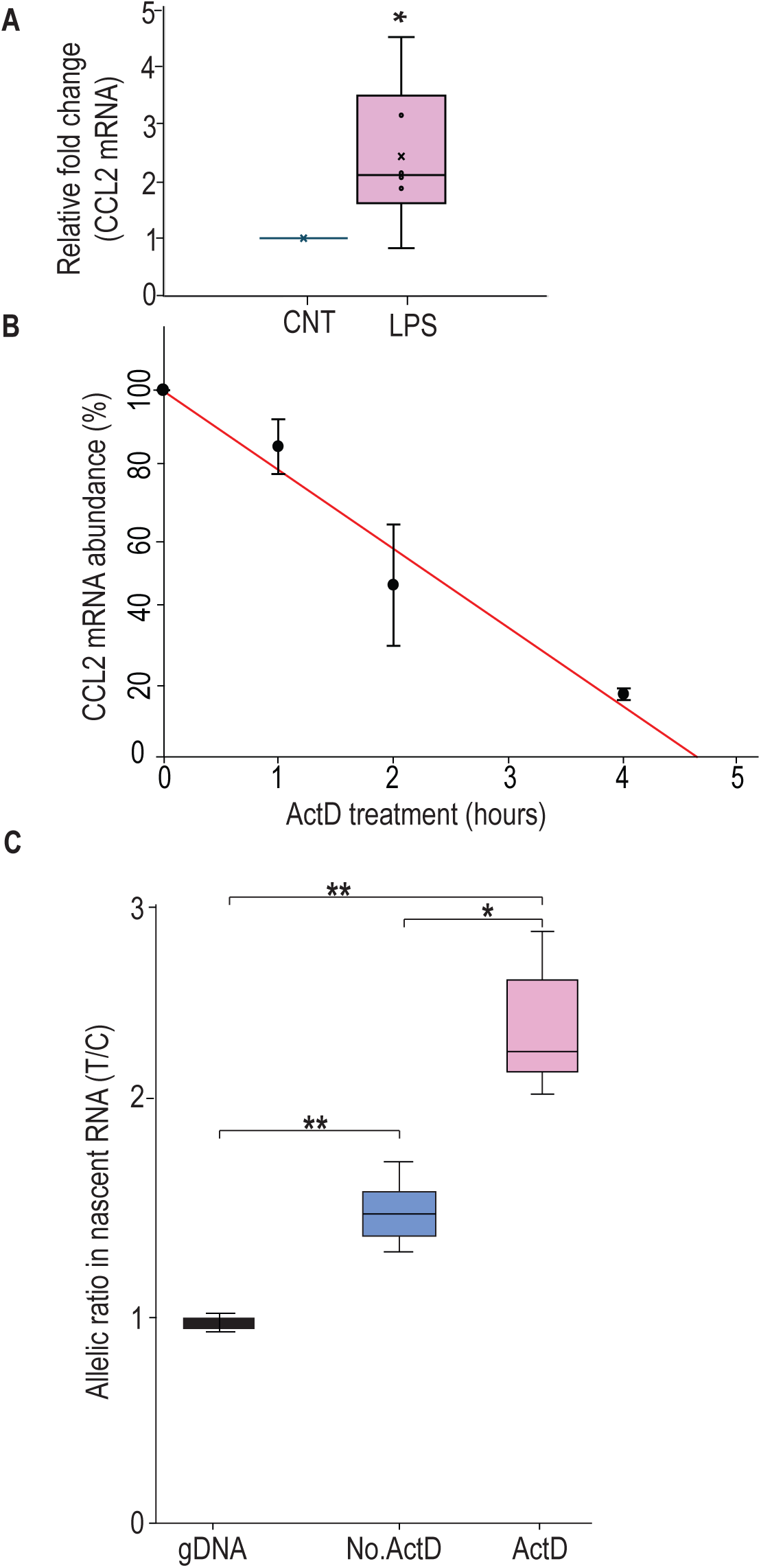
rs13900T confers greater stability to *CCL2* mRNA. **(A)** *CCL2* mRNA expression in peripheral monocytes of heterozygous individuals (n = 6) after treatment with LPS for 3 h and then incubated with 5 µg Act D for indicated times. mRNA was detected by RT-PCR. Results, normalized to 18S rRNA levels, are expressed as folds increase over unstimulated cells (CNT). Levels shown in bar graph represent mean ± SEM of result at time 0 (\**P* = 0.019) versus unstimulated cells (N = 4). **(B)** *CCL2* mRNA half-life, calculated for each condition as the time (in hours) required for the transcript to decrease to 50% of its initial abundance [t ½ = ln (0.5)/slope]. **(C)** Nascent RNA was isolated from treated monocytes from three individuals in the presence and absence of ActD. Allelic ratio was determined after 4 h of incubation with or without ActD. Expression of rs13900 T allele was much higher in ActD-treated samples. The difference between the groups were assessed by ANOVA with Fisher LSD method (\**P* < 0.05, ** *P* < 0.005).

To rule out the confounding effects of preexisting mRNA, the relative stability of rs13900 C-and T-allele bearing transcripts in heterozygous individuals were evaluated using nascent RNA. Using nascent RNA allows accurate determination of mRNA decay by eliminating the effects of preexisting mRNA. Briefly, monocytes were stimulated with LPS in presence of 5-ethynyl uridine (EU) for 3 hours. Cells were then washed and further incubated with or without Actinomycin D for up to 4 hours. Following Actinomycin D treatment, the nascent RNA was captured after 0, 1, 2 or 4 h using click reaction technique. The click reaction adds a biotin handle to nascent RNA which is then captured by streptavidin beads. cDNA was synthesized from the captured nascent RNA, PCR-amplified and expression of the individual alleles was assessed as described above. While the allelic ratio in the no Act-D samples was ∼1.6 in accordance with our initial observation, the allelic ratio was further increased in presence of Act-D. This suggests that post transcriptional mechanisms such as RNA stability may be playing a role in rs13900 mediated CCL2 AEI.

### Bioinformatic analyses of rs13900

While previous experimental studies showed that *CCL2* 3′ UTR binds HuR, it is not known whether rs13900 disrupts or alters HuR binding (Fan et al., 2011; Lebedeva et al., 2011). Therefore, we analyzed the rs13900 flanking region using various bioinformatic software to mine existing whole genome datasets (e.g., PAR-CLIP datasets) and to predict any mRNA structural changes and altered RBP motifs (Supplementary Table 2). We used AURA to examine the colocalization between rs13900 and HuR (Figure 3A). As genetic variants can also alter mRNA secondary structure, we used ViennaRNA package to assessed changes in the *CCL2* secondary structure due to rs13900. As shown in Figure 3B, rs13900 T allele could potentially alter the *CCL2* transcript secondary structure (red arrow). Further bioinformatic analysis was performed using POSTAR3 suite which incorporates HOMER (Heinz et al., 2010) for motif analysis and RNA context (Kazan et al., 2010) that identifies not only known but also predicts relative binding and structural preferences of RBPs. HOMER motif analysis (Heinz et al., 2010) identified a HuR binding motif in the region flanking the rs13900 (Fig. 3C). Figure 3D shows the relative structural preference of HuR to different structural contexts identified by RNAcontext, where the letters P, L, U, M indicate that the nucleotide is paired (P), in a hairpin loop (L), in an unstructured (or external) region (U), or Miscellaneous (M). The M category includes various unpaired contexts such as nucleotide localizing to a bulge, internal loop or multiloop. The rs13900 C allele to rs13900 T allele transition is predicted to form a stem (Figure 3B) which is predicted to increase HuR binding. The score of 1e was obtained using RBP-Var, a bioinformatics tool that scores variants involved in post-transcriptional interaction and regulation (Mao et al., 2016). Here, the annotation system rates the functional confidence of variants from category 1 to 6. While Category 1 is the most significant category and includes variants that are known to be expression quantitative trait loci (eQTLs), likely affecting RBP binding site, RNA secondary structure and expression, category 6 is assigned to minimal possibility to affect RBP binding. Additionally, subcategories provide further annotation ranging from the most informational variants (a) to the least informational variant (e). Reported 1e denotes that the variant has a motif for RBP binding. Although the employed scoring system is hierarchical from 1a to 1e, with decreasing confidence in the variant’s function. However, all the variants in the category 1 are considered potentially functional to some degree. Taken together, our bioinformatic analyses suggested that the rs13900 allele could potentially alter the binding of HuR to *CCL2* 3′ UTR.

**Figure 3.**
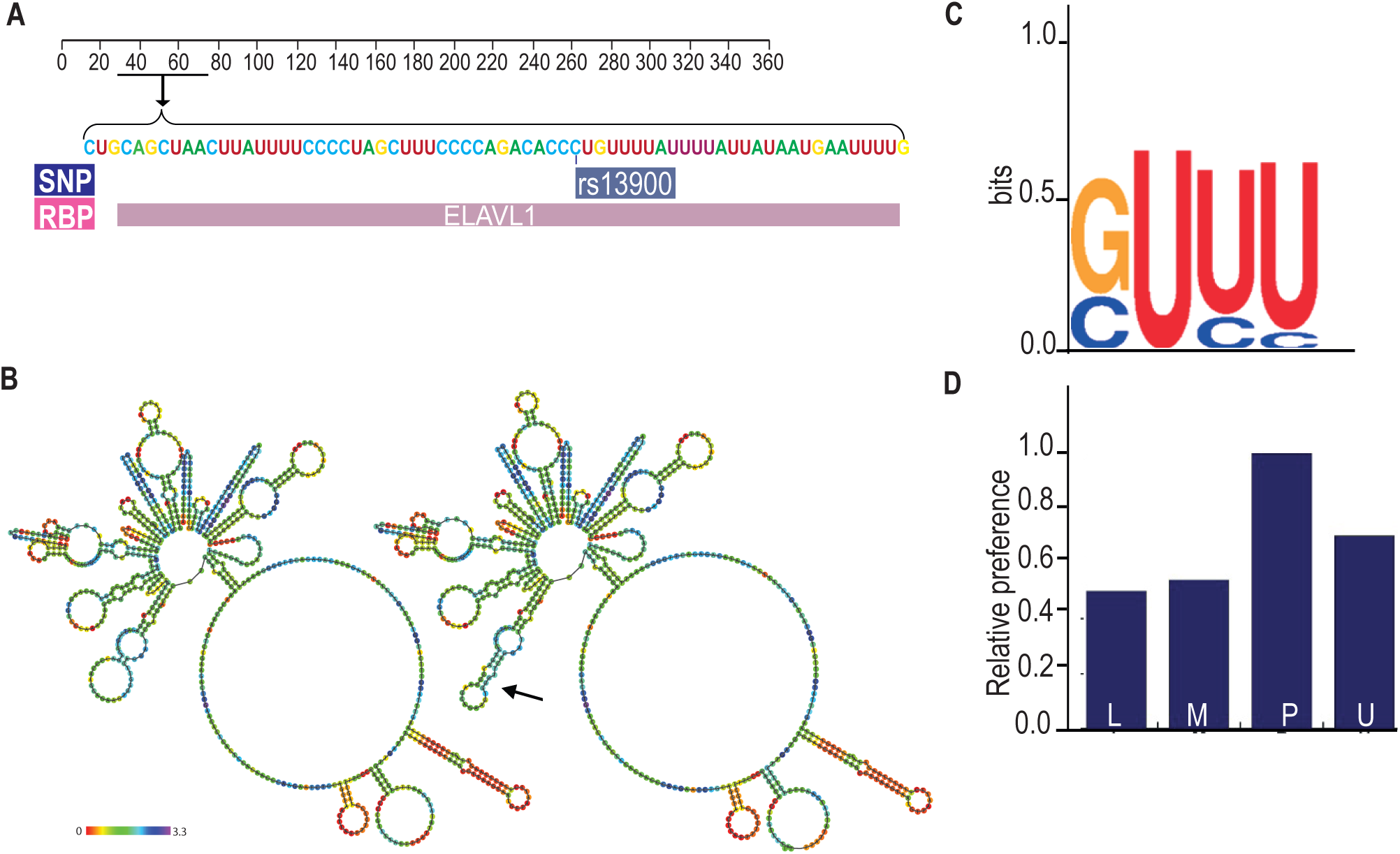
Bioinformatic analysis of the rs13900. **(A)** Validation RBP binding sites and polymorphism located on the 3′ UTR of *CCL2* transcript from the Atlas of UTR Regulatory Activity and analysis of ENCODE genome-wide data sets detected specific enrichment of HuR (ELAVL1) at the region that contains the rs13900. **(B)** Predicted changes in the secondary structure using Vienna RNA package 2.0; changes in secondary structure are indicated by arrows. **(C)** Sequence logo of HuR binding site as determined by HOMER. **(D)** Relative structural preference of HuR to different structural contexts, the letters P, L, U, M indicate that the nucleotide is paired (P), in a hairpin loop (L), in an unstructured (or external) region (U), and Miscellaneous (M) region.

### Differential binding of HuR to rs13900 C and T alleles *in vitro*

To experimentally verify our bioinformatic findings on rs13900, we utilized RNA electrophoretic mobility shift assay (REMSA) to determine whether the region of CCL2 3′ UTR that flanks rs13900 binds *in vitro* to HuR and if there are allelic differences in binding. Purified labeled single-stranded oligoribonucleotides corresponding to either rs13900 C or rs13900 T alleles were incubated with 10 µg of whole cell extracts. We tested whether the bound complexes contained HuR by performing antibody mediated supershift assays. As shown in Figure 4A and 4B, a predominant shift was observed for the oligoribonucleotide corresponding to rs13900 T allele (lane 8). Notably, there was an approximately 7-fold difference (*P* < 0.005) in oligoribonucleotide/HuR/antibody complexes generated with oligoribonucleotide corresponding to rs13900 T allele which compared to those generated with rs13900 C allele (lane 4). The specificity of the complex formation was confirmed by using a non-specific antibody (Figure 4A, lanes 3 & 7). To rule out the possibility that additional RBPs are involved in this complex formation, we used purified HuR protein in mobility shift assays. These studies with purified protein further confirmed that oligoribonucleotides corresponding to rs13900 C or T alleles differentially bound HuR (Figure 4C). Taken together, these studies provided experimental validation that the rs13900 may influence differences in binding affinity to HuR (Figure 4D).

**Figure 4.**
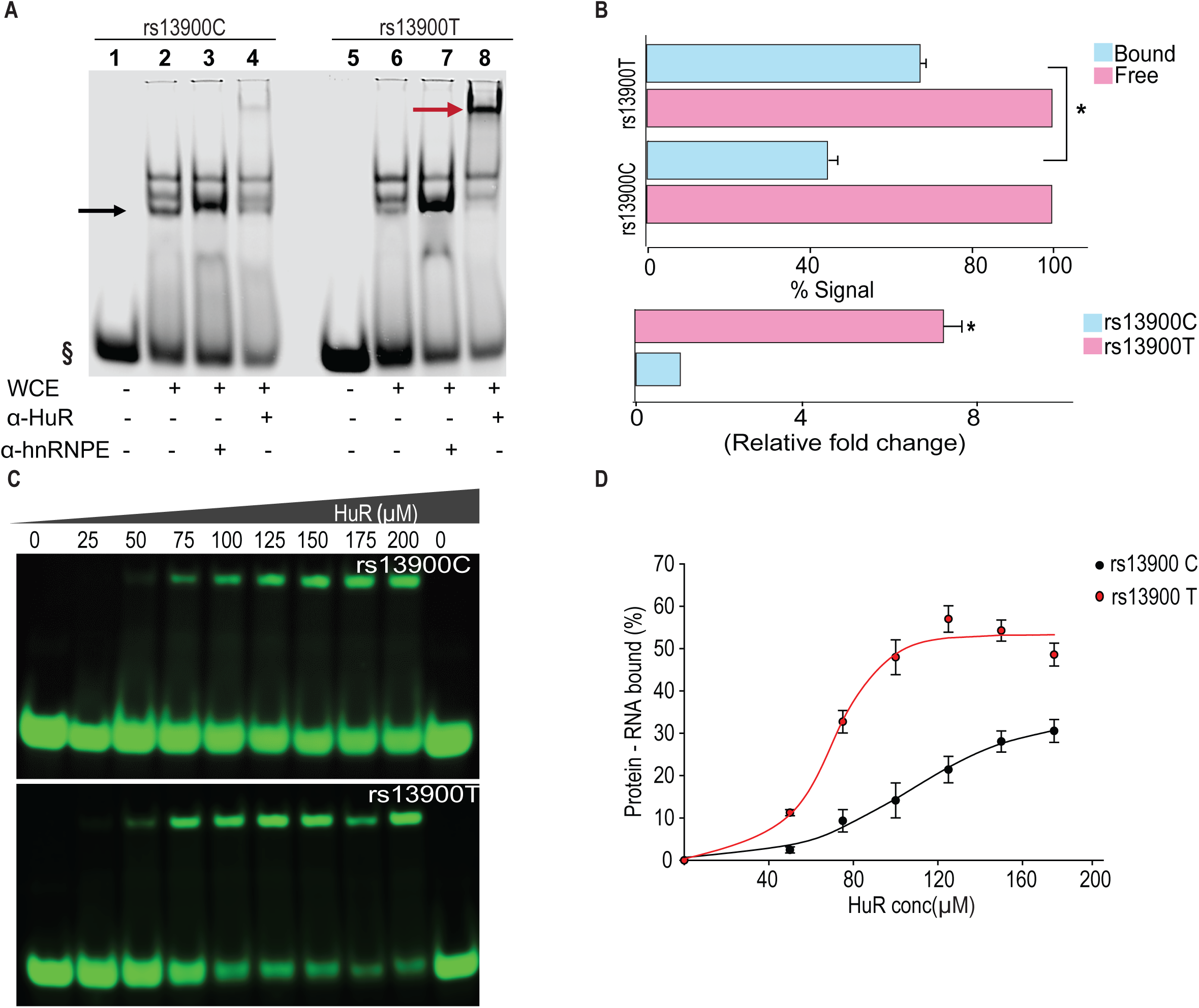
The rs13900 T allele shows increased in vitro binding of HuR. **(A)** REMSA with labelled oligoribonucleotide containing either rs13900 C or T allele and whole cell extracts from 293T cells. § denotes free probe; black arrow, bound probe; red arrow, supershift. **(B)** Representative quantitative densitometric analysis of the antibody shifted complexes suggested increased HuR binding to the oligoribonucleotide bearing rs13900 T allele. The signals in the bound fraction(s) were normalized using the free probe (N = 4). The top panel represents the data from four independent experiments (mean ± SEM). Statistical analyses were performed using Student’s t test (\**P* < 0.001). The bottom panel shows the relative fold enrichment of the bound protein complexes to the oligoribonucleotide containing the rs13900 T allele relative to that containing the rs13900 C allele . Statistical significance was calculated using Student t test (\**P* < 0.001) **(C)** REMSA with labelled oligoribonucleotides containing either rs13900 T or C allele and purified HuR protein. **(D)** Plot showing the fraction of bound rs13900 C or rs13900 T oligoribonucleotides with increasing HuR concentration.

### Differential binding of HuR to rs13900 C and T alleles *ex vivo*

We next tested the hypothesis that the rs13900 is associated with altered binding affinity to HuR *ex vivo*. For this, we performed RNA immunoprecipitation in monocyte/macrophages derived from four heterozygous individuals. Immunoprecipitations were performed using cytoplasmic lysates of macrophages treated with LPS using an affinity purified HuR antibody or IgG. Figure 5A shows the relative enrichment of HuR in the immunoprecipitated fraction compared to the IgG control. The HuR immunoprecipitated fraction showed significant enrichment (*P* < 0.05) of the region encompassing rs13900 when compared to the IgG control (Figure 5B-C). *CCL2* transcript was enriched ∼10-fold in the HuR immunoprecipitated fraction in comparison to the IgG bound fraction (Figure 5C). Notably, transcripts corresponding to rs13900 T allele were enriched relative to transcripts containing rs13900 C allele (*P* < 0.05) in the anti-HuR RIP complexes (Figure 5D).

**Figure 5.**
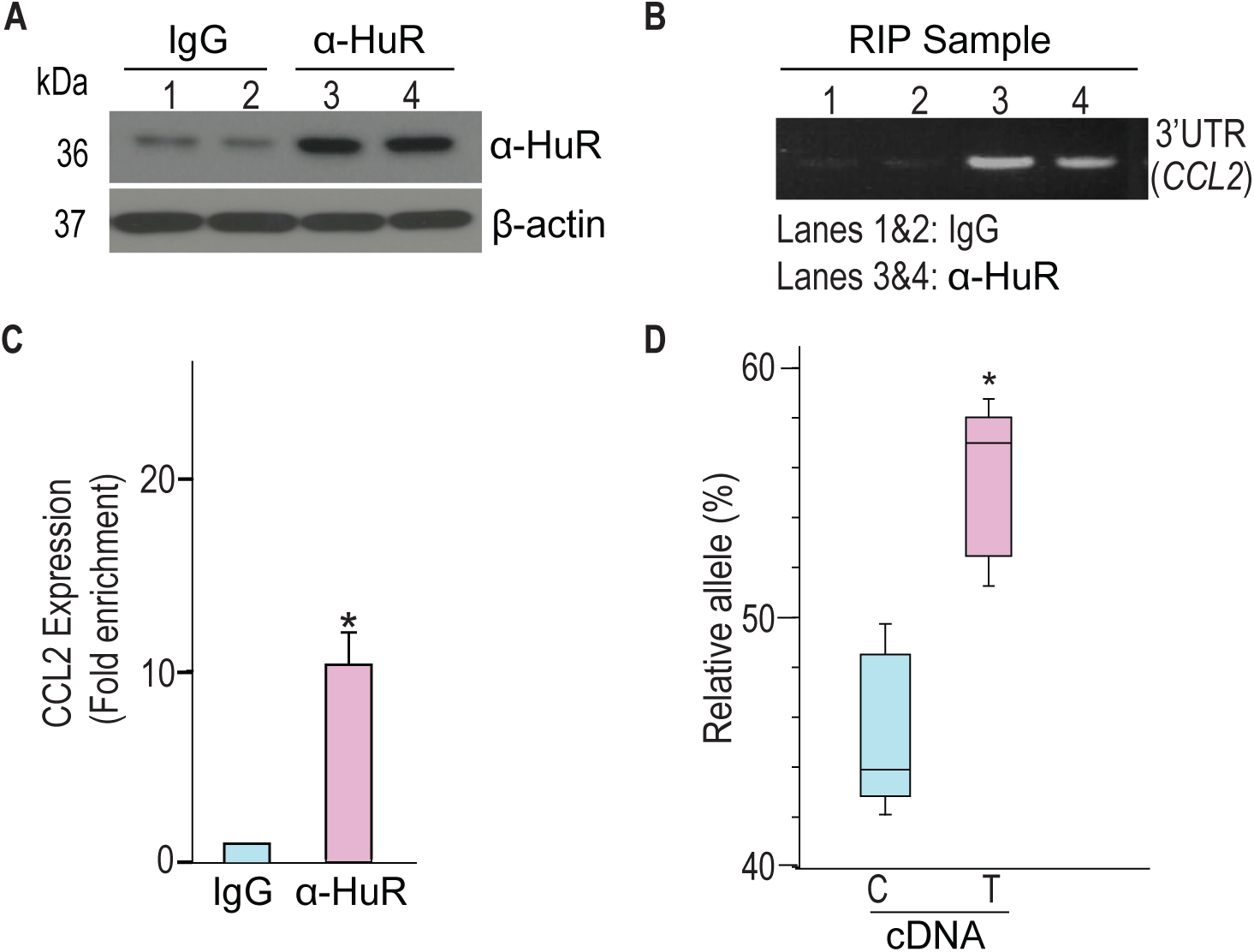
**rs13900 C and T alleles are associated with differential binding to HuR ex vivo**. **(A)** HuR enrichment in immunoprecipitated material from macrophages stimulated with LPS. **(B)** *CCL2* 3′ UTR was detected at significant levels in the samples precipitated by α-HuR antibody when compared to the control IgG. **(C)** *CCL2* mRNA expression in anti-HuR antibody enriched immunoprecipitated material analyzed by RT-qPCR (N = 4). Statistical significance was calculated using Student t-test (\**P* < 0.005). The error bars represent SEM. **(D)** Relative expression levels for rs13900 C and T alleles in macrophages stimulated with LPS (N = 4). Statistical significance was calculated using a Student t-test (\**P* < 0.005).

### rs13900 T allele confers increased mRNA stability in reporter assays

As HuR is implicated in mRNA stability of many mRNA transcripts including *CCL2*, we tested the hypothesis that *CCL2* 3′ UTR influences its stability and that rs13900 modulates this effect. We constructed reporter plasmids that harbor the *CCL2* 3′ UTR containing either rs13900 C or rs13900 T allele (Figure 6A) and nucleofected them into HEK293 cells as described in Materials and Methods. Luciferase activity was measured after 24 h post-transfection (Figure 6B). Our results indicated that the presence of *CCL2* 3′ UTR significantly (*P* < 0.05) reduced the luciferase activity. However, cells transfected with plasmids bearing the rs13900 T allele showed higher luciferase activity when compared with cells transfected with plasmids bearing the rs13900 C allele, suggesting that the presence of rs13900 T allele conferred increased stability to the transcript (*P*<0.05). We next analyzed the influence of HuR overexpression in reporter vectors containing *CCL2* 3′ UTR with either rs13900 C or T alleles. Overexpression of HuR led to a significant increase in luciferase activity of the reporter vector bearing rs13900 T allele (*P* < 0.05), However, HuR overexpression had no significant effect on the luciferase activity of the reporter vector bearing the rs13900 C allele (Figure 6C). Conversely, we examined the effect of HuR on the expression of the reporter assay by co-transfection of HuR siRNA and luciferase reporter constructs. While HuR knockdown had no effect on luciferase activity of the reporter construct bearing rs13900 C allele, it caused a significant reduction in luciferase activity of construct bearing the rs13900 T allele (*P* < 0.05) (Figure 6D). The overexpression and knockdown of the HuR in the nucleofected cells was confirmed using Western blots (Figure S3).

**Figure 6.**
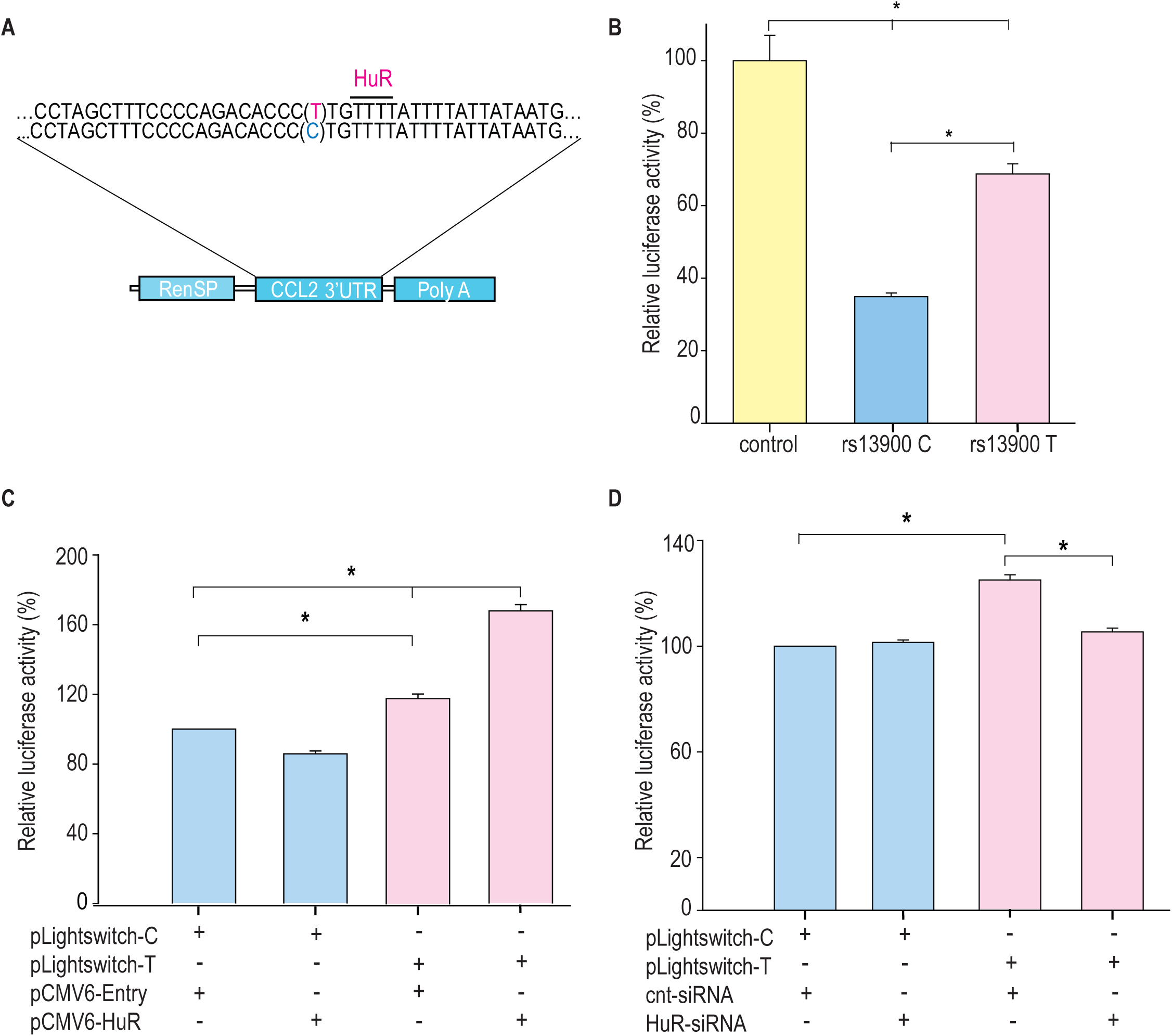
Differential effects of rs13900 alleles in reporter assays and role of HuR. **(A)** Schematic representation of the luciferase reporter vectors containing *CCL2* 3’ UTR with either rs13900 C or T allele. **(B)** HEK-293 cells were transfected with the equal quantities of *CCL2* 3′ UTR reporter vectors and luciferase activity was measured 48 hrs later. The relative luciferase activities of the 3′ UTR reporter plasmids were expressed as percent reduction in the luminescence when compared to the control vector that was set to 100% after normalizing for the protein content of the lysates. The error bars indicate the standard error of mean and statistical significance was calculated using two-tailed Student’s *t* test (\**P* < 0.05). **(C)** HEK-293T cells were transfected with either pCMV6-HuR (0.5 μg) or pCMV-Entry (0.5 μg) and after 72 hours they were co-transfected with the two plasmid constructs (0.5 μg). Twenty-four hours after transfection the relative change in luciferase activity was determined. **(D)** Cells were co-transfected with 125 pmol HuR siRNA or control siRNA and with the two plasmid constructs (0.5 μg). Twenty-four hours after transfection the relative change in luciferase activity was determined (normalized to total protein concentration, data from four independent experiments (mean ± SEM; N = 4). Statistical analyses were performed using Fisher LSD method (\**P* < 0.05).

### Role of HuR-rs13900 interactions in *CCL2* mRNA translatability

The differential interactions with HuR by rs13900 C and rs13900 T alleles could potentially alter *CCL2* mRNA translatability as HuR could increase mRNA translation (Schultz et al., 2020). Therefore, we assessed the relative allelic enrichment in the monosomal and polysomal fractions obtained from MDMs from two individuals exhibiting AEI. Translationally active and inactive pools of RNA were fractionated by isolating the monosomal and polysomal fractions from MDMs on sucrose gradients after the cells were treated with cycloheximide to block translation. The distribution of *CCL2* mRNA in macrophages cultured in presence or absence of LPS for 3 h is depicted in Figure S4. As previously reported by others, LPS treatment resulted in a distinct shift of the mRNAs from monosomal fraction to the polysomal fraction (Schott et al., 2014). Consistent with this prior report, 60.4% of the *CCL2* mRNA was associated with polysomal fraction following stimulation of cells with LPS. We assessed the differences in allelic enrichment in cytosolic, monosomal, and polysomic fractions and found that the polysomal fractions showed the enrichment of the rs13900 T allele (Supplementary Table 3). However, this differential loading of rs13900 T allele was noted for one donor in cytosolic and monosomal fractions as well.

To further address this question, we used a reporter-based system to assess the effect of rs13900 C and T alleles on the translatability as previously reported (Zhang et al., 2017). To measure translatability, luciferase mRNA and protein were measured simultaneously, and translatability was calculated as luciferase activity normalized by the luciferase mRNA levels after adjusting for protein and 18S rRNA (Figure 7A-C). Our results suggest that the rs13900 may alter the mRNA translatability in addition to the transcript stability.

**Figure 7.**
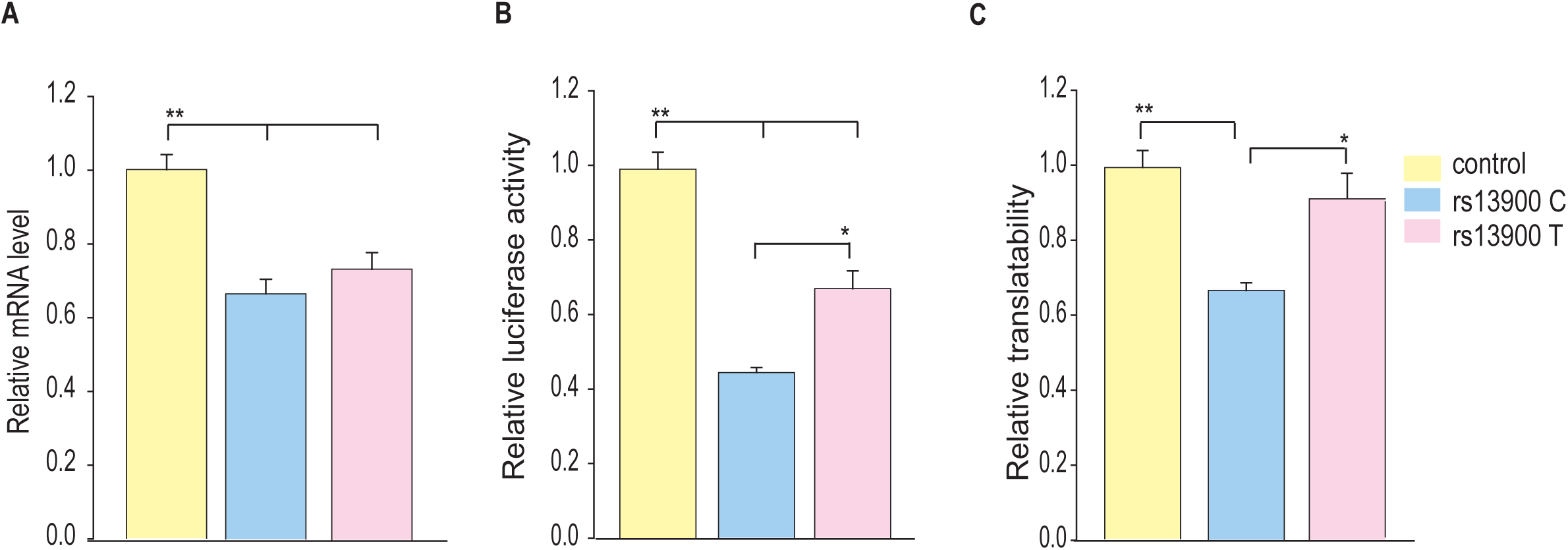
Influence of rs13900 C and T alleles on *CCL2* translatability. **(A)** HEK-293 cells were transfected by nucleofection with control and *CCL2* 3′ UTR reporter constructs with rs13900 C and T alleles. The nucleofected cells were plated separately and harvested for total RNA isolation or lysed for mRNA level or protein level expression of luciferase respectively after 24 h. The reporter mRNA levels from the transfected 293 T cells were quantified by qRT-PCR, and 18S rRNA was used for normalization. **(B)** The relative luciferase activities of the 3′ UTR reporter plasmids were expressed as a percentage reduction in the luminescence when compared to the control vector that was set to 100% after normalizing for the protein content of the lysates. **(C)** mRNA translatability was calculated as luciferase activity normalized by the reporter luciferase mRNA level. The error bars indicate the standard error of mean from six independent experiments (N = 6) and statistical significance was calculated using ANOVA and post hoc contrast with Fisher LSD method. (\**P* < 0.05, \*\**P* < 0.005).

### Differential effect of HuR over-expression on the *CCL2* rs13900 T allele

Our *in vitro* and *ex vivo* data indicate that HuR positively regulates *CCL2* transcript stability by increased binding to the T allele. To determine a direct functional relationship between HuR and *CCL2* AEI, we used a lentiviral overexpression system. We either transduced HuR specific (pCMV6-HuR) or non-specific control shRNAs (CMV-null) into the monocytes obtained from donors who were either homozygous for rs13900 C or T allele. Ready to use GFP-tagged pCMV6-HuR or CMV-null lentiviral particles were transduced into macrophages in the presence of polybrene at an MOI of 1. Cells were processed 72 h following virus addition for the analysis of HuR and *CCL2* mRNA expression. Using this lentiviral system, both high transduction efficiency (90%) and high expression levels were achieved in primary human macrophages (Figure S5). Compared with CMV-Null particles, substantial overexpression of HuR was noted with pCMV6-HuR lentiviral particles (Figure 8A). HuR overexpression was associated with a higher expression of *CCL2* mRNA in persons homozygous for T allele relative to those who were homozygous for C allele (Figure 8B).

**Figure 8.**
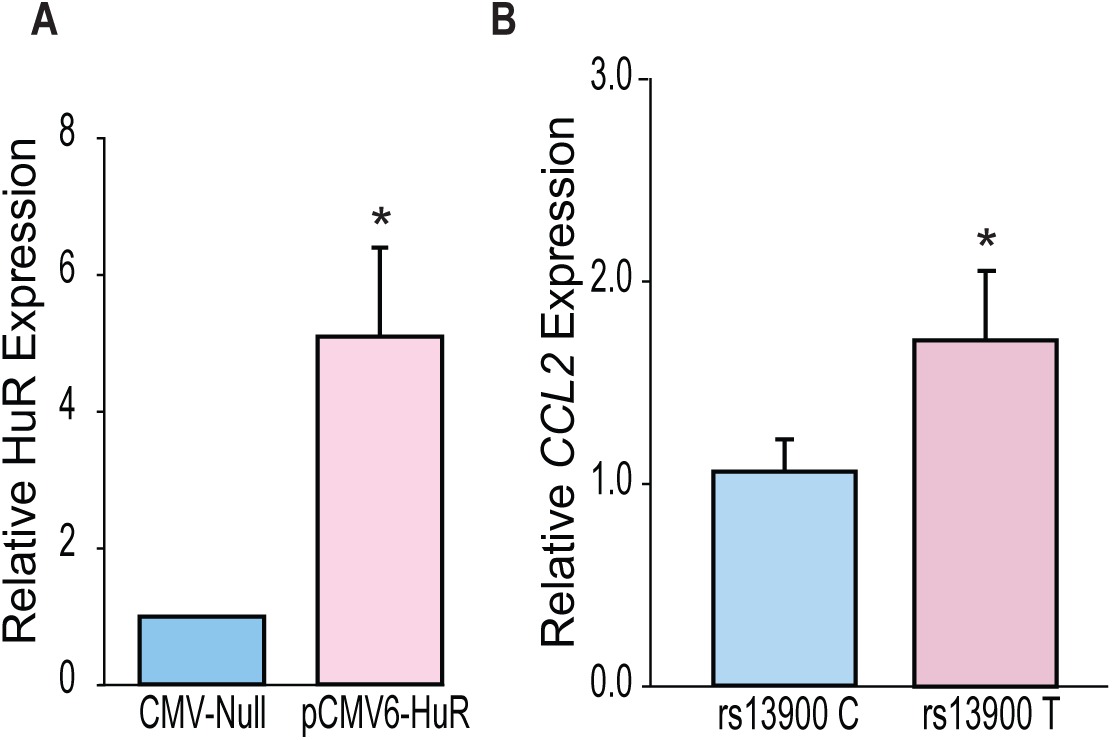
HuR differentially regulates *CCL2* haplotypes. **(A)** HuR expression in primary human macrophages following lentiviral transduction. Macrophages obtained from four individuals who are homozygous for either rs13900 C or rs13900 T allele were transduced with either CMV-null or HuR expressing lentiviral particles (pCMV6-HuR) for 72 hours followed by LPS stimulation for 3 h. **(B)** *CCL2* expression determined by RT-qPCR. Error bars represent SEM. Statistical analyses were performed using Student’s t test (* *P* < 0.05).

## Discussion

Genome-wide association studies (GWASs) have led to the discovery of numerous disease susceptibility genetic variants/genes and biological pathways involved in specific diseases, including many with immunological etiologies (Buniello et al., 2019; Manolio, 2010; Visscher et al., 2017). Most of the disease-associated genetic variants identified are present in the non-coding regions of the genome (Maurano et al., 2012; Zhang and Lupski, 2015; Zou et al., 2019). However, characterizing the functional consequence of these genetic variants in the regulatory regions remains a significant challenge to date. About 3.7% of the non-coding variants localize to the untranslated regions (Steri et al., 2018), suggesting post-transcriptional mechanisms such as mRNA stability and translatability could determine disease susceptibility (Maurano et al., 2012).

The *CCL2* locus exemplifies the difficulty in dissecting the functional consequences of cis-regulatory and non-coding genetic variants. Our analysis of regulatory elements in an ∼20 kb region upstream of the human *CCL2* coding region identified highly conserved enhancers and other regulatory elements in the *CCL2* locus (Bonello et al., 2011). We identified SNPs with strong LD in this “super-enhancer” and determined their impact on transcriptional activity, which was minimal (Pham et al., 2012). For example, we tested the role of rs7210316 and rs9889296, which had an r^2^ > 0.9 with rs1024611 in CEU population and were located ∼6.3 kb and ∼9.2 kb upstream of it (Pham et al., 2012). The genetic regions that encompass and flank these two SNPs either had no or minimal transcriptional activity and lacked the epigenetic markers that have been traditionally associated with gene activation. Additionally, conflicting context-dependent results have been obtained for the transcriptional activities associated with reporter vectors containing rs1024611 A and G alleles, which may have been due to the use of different lengths of the *CCL2* cis-regulatory regions employed in the transcriptional assays. To avoid such context-dependent confounding, we used chromatin annotation, a powerful approach to identify functional SNPs, to generate reporter constructs that span a 6 kb cis-regulatory region (Pham et al., 2012). This approach allowed us to directly compare the roles of four correlated SNPs (rs1860190, rs2857654, rs1024611, and rs2857656) on *CCL2* transcriptional activity. We found no significant differences in transcription strength between these two constructs when transfected into primary human astrocytes (Pham et al., 2012). Furthermore, to understand the basis for increased CCL2 expression associated with the rs1024611 G allele, we and others have demonstrated that several transcription factors bind differentially to *CCL2* polymorphisms, including IRF-1, PARP-1, STAT-1, Prep1/Pbx complexes (Gonzalez et al., 2002; Mummidi, Bonello and Ahuja, 2009; Page et al., 2011; Wright et al., 2008). However, these previous studies could not unequivocally demonstrate the mechanistic link between differential binding of transcription factors and transcriptional activity and the physiological relevance of implicated transcription factors in CCL2 expression. Our comprehensive studies outlined here demonstrate that rs13900 T allele differentially binds to the RBP HuR, which could alter CCL2 mRNA stability and gene expression. Furthermore, we provide multiple lines of functional evidence for rs13900/HuR role in *CCL2* AEI, including reporter vector assays and HuR overexpression and depletion experiments.

Our previous finding that rs13900 is in perfect LD with rs1024611 has allowed us to exploit the powerful AEI technique and mechanistically link the *CCL2* rs1024611G-rs13900T haplotype to increased expression (Pham et al., 2012). These findings, together with previous knowledge that *CCL2* is subjected to post-transcriptional regulation (Hao and Baltimore, 2009) and regulated by RBP (Fan et al., 2011), raised the possibility that rs13900 could be functional and impact mRNA stability and translatability. In this study, we determined the half-life of *CCL2* to be 1.76 h, reinforcing the fact that post-transcriptional regulation is a key feature of *CCL2* expression (Fig.2B). Since transcription and RNA degradation are tightly linked, and transcriptional inhibition may lead to mRNA decay, we assessed the allelic differences in expression in nascent RNA obtained from heterozygous individuals. We found that the *CCL2* transcripts with rs13900 T allele have increased stability relative to those with rs13900 C allele (Figure 2C). In addition, both rs1024611 and rs13900 have suggestive associations with *CCL2* expression in *cis*-expression quantitative trait loci (eQTL) analyses. For example, rs1024611 and rs13900 are associated with CCL2 expression in MDM infected with *Listeria monocytogenes* (Nedelec et al., 2016) (−log10(p) = 3.19; Effect size = 0.29±0.084) (Kwong et al., 2021). Remarkably, rs13900 is also identified as a cis-splicing quantitative trait locus (cis-sQTL) for *CCL2* in both untreated and treated monocytes (Alasoo et al., 2018; Kwong et al., 2021; Quach et al., 2016). Recent evidence suggests that genetic variation influencing RNA splicing could play an important role in determining complex phenotypic traits (Li et al., 2016). While rs13900 serves as an sQTL for several *CCL2* transcripts variants, we provide an illustrative example here. rs13900 showed significant associations with the canonical *CCL2* transcript, ENST00000225831 in LPS treated monocytes (−log10(p) = 9.88; Effect size = 0.57±0.084) and with a CCL2 transcript ENST00000580907 (−log10(p) = 9.88; Effect size = −0.57±0.084) that encodes a truncated CCL2 protein (Kwong et al., 2021). While the importance of these associations needs more in-depth studies, our observation that *CCL2* transcripts with rs13900 T allele have a slower degradation could explain the increased CCL2 expression in individuals with rs1024611G-rs13900 haplotype as modest changes in mRNA stability can lead to significant effects on mRNA and protein abundance.

Post-transcriptional gene expression is regulated through interactions between the *cis*-elements in mRNAs and their cognate RBPs (Wu and Brewer, 2012). Studies indicate presence of post-transcriptional operons or regulons – unique subsets of RNAs that associate with RBPs, which coordinate their localization, translation, and degradation. Our bioinformatics analysis and experimental data confirm HuR binds to the region spanning the rs13900. iClip data from HeLa cells (Wang et al., 2010), suggested that the RBPs TIAL1 (T cell intracellular antigen-1 like protein) and U2AF65 (U2 snRNP auxiliary factor large subunit) could also bind to this region of the *CCL2* transcript (Mao et al., 2016). TIAL1 has been proposed to act as a cellular sensor and has been associated with a transcriptome associated with control of inflammation, cell-cell signaling, immune-suppression, angiogenesis, metabolism and cell proliferation (Reyes, Alcalde and Izquierdo, 2009). The role of TIAL1 in *CCL2* expression is not known and additional studies are needed to determine if its binding contributes to *CCL2* AEI. U2AF65 is a widely expressed splicing factor that associates with RNA polymerase II to bind upstream 3′ splice sites to facilitate splice site pairing in higher eukaryotes (Hollander et al., 2016). In addition, a change in RNA structure due to a SNP could indirectly alter accessibility of additional regions to RBPs and further studies are required to determine any such changes (Shatoff and Bundschuh, 2020).

The Hu family contains 4 members, of which HuR is ubiquitously distributed. HuR consists of three RNA recognition motifs (RRMs) that are highly conserved and canonical in nature (Ripin et al., 2019). In absence of RNA the three RRMs are flexibly linked but upon RNA binding they transition to a more compact arrangement. Mutational analysis revealed that HuR function is inseparably linked to RRM3 dimerization and RNA binding. Dimerization enables recognition of tandem AREs by dimeric HuR (Pabis et al., 2019) and explains how this versatile protein family can regulate numerous targets found in pre-mRNAs, mature mRNAs, miRNAs and long noncoding RNAs. HuR shuttles between the nucleus and cytoplasm and is likely involved in the transport and stabilization of mRNA. HuR action is antagonized by RNA destabilizing proteins such as TTP (tristetraprolin) and AUF (ARE/poly(U)-binding/degradation factor 1) (Wu and Brewer, 2012). The role of HuR in chemokine gene regulation has been extensively studied in human airway epithelium cells (Fan et al., 2011), they identified that *CCL2* is one of the top targets for HuR and demonstrated that HuR associated with *CCL2* 3′ UTR *in vitro* and *CCL2* expression could be modulated by changes in HuR levels using overexpression and RNAi-based experiments. Also, HuR levels influenced *CCL2* mRNA decay. In a mouse model of alcoholic liver disease, HuR was found to play a key role in NOX4 mediated increase in *CCL2* mRNA stability (Sasaki et al., 2017). Notably, HuR has also been implicated in mRNA stability of CX3CL1 (Matsumiya et al., 2010) and IL-8 (Choi et al., 2009) among others. Another study showed that CCL2 itself can induce nuclear to cytoplasmic translocation of HuR and stabilize vascular endothelial growth factor-A (*VEGFA*) mRNA in CD14^+^CD16^low^ inflammatory monocytes, thus raising the possibility that CCL2 may stabilize its own message in an autocrine fashion (Morrison et al., 2014). These studies further bolster a role for HuR in *CCL2* expression and support our finding that differential binding to HuR alters its expression level.

While there are several examples of genetic variants that influence mRNA stability (Akdeli et al., 2014; Duan et al., 2013; Vilmundarson et al., 2021; Wang, Pitarque and Ingelman-Sundberg, 2006), very few studies have examined the role of RBPs such as HuR in differentially influencing gene expression levels in humans (Steri et al., 2018). A notable exception is the report showing that RBP AUF1 regulates the allele-specific stability of thymidylate synthase (Pullmann et al., 2006). Another study showed that a type 2 diabetes associated polymorphism in the 3′ UTR of *PPP1R3* alters distance between two ARE motifs and results in differential binding of protein complexes and may be associated with altered mRNA stability (Xia, Bogardus and Prochazka, 1999). Vilmundarson et al. demonstrated that HuR differentially modulates *IRF2BP2* translation through a 3′ UTR polymorphism that is associated with increased coronary artery calcification (Vilmundarson et al., 2021). Our exploratory studies did not resolve whether the rs13900 allelic variation leads to differences in polysomal loading and additional studies are required to address this issue. Another important mechanism by which disease associated genetic variants in the UTRs could influence mRNA stability and translatability is by disrupting or creating microRNA binding sites (Huang et al., 2019). We examined the region flanking the rs13900 and found that there are several predicted binding sites for miRNAs. Among these, several miRNAs have minimum free energy ≤ –25 kcal/mol suggesting that they can potentially play a role in *CCL2* expression and will be investigated in future studies. Of note, a recent study showed that RBPs may play an important role in miRNA mediated gene regulation (Kim et al., 2021).

RBPs regulate the protein expression from a given mRNA by modulation of its half-life, subcellular localization, and ribosomal recruitment (Glisovic et al., 2008). However, mRNA abundance and stability are not always predictive of protein synthesis: relative mRNA abundance/stability and translation levels of a given gene can vary significantly and determined by an assortment of post-transcriptional events (Liu, Beyer and Aebersold, 2016). Our previous findings and results from the present study suggest that there is a close association between *CCL2* rs1024611-rs13900 haplotype, mRNA, and protein expression. Nevertheless, we cannot completely rule out the contribution of transcription-based mechanisms to the allelic expression imbalance at the *CCL2* locus. Previous studies have suggested that there is a high level of conservation for interactions between RBPs and their target molecules and that RNA-mediated gene regulation is less evolvable than transcriptional regulation (Payne, Khalid and Wagner, 2018). Thus, it is remarkable that the rs13900 alters HuR binding and impacts both transcript stability and translatability. Our bioinformatic analysis shows that rs13900 could potentially lead to dramatic changes in the secondary structure of *CCL2* mRNA (Fig. 3B). A recent study using bioinformatic analyses has shown that SNPs may affect RNA-protein interactions from outside binding motifs through altered RNA secondary structure (Shatoff and Bundschuh, 2020). We cannot rule out the possibility that HuR may differentially bind to other regions of *CCL2* transcript due to changes in secondary structure and therefore needs to be examined in future studies.

Given our findings that rs13900 modulates HuR binding and thereby influences *CCL2* expression, targeting HuR-ARE interactions could offer a promising therapeutic avenue for conditions involving heightened monocyte/macrophage recruitment, such as inflammatory, cardiovascular, and neoplastic diseases. Indeed, existing small-molecule inhibitors of HuR (e.g., MS-444, KH-3, and CMLD-2) (Chaudhary et al., 2023; Fattahi et al., 2022; Lang et al., 2017; Liu et al., 2020; Wang et al., 2019; Wei et al., 2024) highlight the feasibility of disrupting HuR function in vivo, suggesting that co-targeting HuR and the rs13900 may represent a novel precision-based strategy for monocyte/macrophage-driven disorders.

Both rs1024611 and rs13900 are in high LD with rs3091315, which is detected in the GWA studies of CD and IBD (de Lange et al., 2017; Franke et al., 2010; Kenny et al., 2012; Liu et al., 2015; Palmieri et al., 2010). rs3091315 is classified as an upstream risk variant and is located in the intergenic region between *CCL2* and *CCL7*. The GWA risk allele for CD is rs3091315-A which is correlated with rs1024611(A) and rs13900(C) alleles. While the biological role of *CCL2* in CD and IBD are well recognized (Darkoh et al., 2014; Maharshak et al., 2010; Martin et al., 2019), the results from genetic association studies are not consistent and the disease outcomes could potentially differ by population studied (Chen et al., 2016).

In conclusion, our study shows that the disease associations mediated by the *CCL2* rs1024611-rs13900 haplotype may be due to altered mRNA stability mediated through differential binding of an RBP. Given the importance of mRNA stability in immune homeostasis such mechanisms could play a critical role in inter-individual differences in disease pathogenesis.

## Materials and Methods

### Recruitment of study participants, primary cell culture, and genotyping

All research involving human subjects were approved by the Institutional Review Boards (IRBs) of the University of Texas Health San Antonio, San Antonio, Texas and University of Texas Rio Grande Valley, Edinburg, Texas and Texas A&M University-San Antonio. Written, informed consent from each individual for participation in our study was obtained following the IRB approval. A total of 47 unrelated individuals were recruited into the study. Data and samples from the study participants were obtained at the First Outpatient Research Unit (FORU), UT Health San Antonio, San Antonio, Texas. The rs13900 polymorphism was detected by TaqMan predesigned SNP genotyping assay (Applied Biosystems, CA, USA, Cat. No. 4351379). Briefly, genomic DNA samples were obtained from peripheral blood of healthy study participants using QIAamp DNA Blood Mini Kits (QIAGEN, Cat. No 51104) in accordance with the manufacturer’s protocol, checked for quality and concentration, and stored at −80°C until used. Genotyping of rs13900 was performed on 10 ng of genomic DNA using TaqMan genotyping Master mix in a 10 µL reaction volume. PCR was performed on the Quantstudio 12K Flex (Applied Biosystems) in 384-well plates. The temperature cycling conditions consisted of an initial enzyme heat-activation step of 10 min at 95°C and 40 cycles of a 3-step amplification profile of 20 sec at 95°C for denaturation, 1 min at 60°C for annealing, and 30 sec for 72°C for extension. QS12K software (Applied Biosystems, CA, USA). was used to score the alleles. Whole blood samples were collected from study participants who were either homozygous (either C/C or T/T) or heterozygous (C/T) for rs13900 at the initial screening for the genotype status, followed by recalling selected individuals for detailed studies. Peripheral blood mononuclear cells (PBMC) were isolated by a Ficoll density gradient centrifugation. Total genomic DNA and RNA were isolated from 0.5x10^6^ PBMC following stimulation with 1 µg/mL LPS (Millipore Sigma, Cat No. L2630) for 3 h as described previously (Pham et al., 2012; Sharif et al., 2007). CD14+ monocytes were purified using EasySep Human Monocytes Isolation Kit (negative selection kit; Stem cell Technologies, Cat No. 19359) according to manufacturer’s instructions. Monocytes were treated with 50 ng/ mL M-CSF (Peprotech, Cat No. 300-25) for 72 h to induce macrophage differentiation, and flow cytometric measurement of surface markers CD64 (BD-Pharmigen, Cat No. 555522), CD206 (BD-Pharmigen, Cat No.564060), and CD44 (BD-Pharmigen, Cat No. 555479) was used to confirm the differentiation (Figure. S6).

### Real-time quantitative PCR (RT-qPCR) assays

Total RNA was isolated using the RNeasy plus mini kit (Qiagen, Cat No. 74134) according to manufacturer’s instructions. 10 μL of total RNA sample (500 ng) was converted to cDNA using MultiScribe Reverse Transcriptase (Applied Biosystems, Cat No. 4311235) with random primers. Real-time quantitative PCR was performed using TaqMan gene expression assays (ThermoFisher Scientific, Cat No. 4331182). Relative expression levels were calculated by applying the 2^−ΔΔCt^ method using 18S rRNA as a reference.

### Capture of nascent RNA

Newly synthesized RNA was isolated using the Click-It Nascent RNA Capture Kit (Invitrogen, Cat No: C10365) following the manufacturer’s protocol. Peripheral blood mononuclear cells (PBMCs) or monocyte-derived macrophages (MDMs) obtained from heterozygous individuals were stimulated with 1 µg/mL lipopolysaccharide (LPS) for 3 hours, followed by a 3-hour pulse with 0.2 mM 5-ethynyl uridine(Jao and Salic, 2008; Paulsen et al., 2013). After the pulse, the culture medium was replaced with fresh growth medium devoid of EU. To assess RNA stability, cells were left untreated or were treated with actinomycin-D (5 µg/mL) and samples were collected at 0, 1, 2, and 4 h post-treatment. The EU RNA was subjected to a click reaction that adds a biotin handle which was then captured by streptavidin beads. The captured RNA was used for cDNA synthesis (Superscript Vilo kit, Cat No: 11754250), PCR amplification, and allelic quantification.

### AEI assessment of *CCL2*

Total RNA and genomic DNA were isolated from primary immune cells (PBMC and macrophages) after they were treated with LPS for 3 h. Total RNA was reverse transcribed as described above. The region encompassing rs13900 was amplified by PCR in a 50 µL reaction mixture containing 2 µL of cDNA or genomic DNA, 25 µL AmpliTaq Gold 360 Master Mix (ThermoFisher Scientific, Cat No: 4398881) and 10 µM each of the forward and reverse primers. The nucleotide sequence of the forward primer and reverse primers were 5′*-*ACCTGGACAAGCAAACCCAA*-*3′ and 5′*-* ACCCTCAAAACATCCCAGGG*-*3′. The following temperature conditions were used: initial denaturation at 95°C for 10 min followed by 40 cycles of denaturation at 95°C for 30 s, annealing at 53.6°C for 30 s, extension at 72°C for 30 s, and final extension at 72°C for 7 min. The amplicons were purified with The GeneJET PCR Purification Kit (ThermoFisher Scientific, Cat No: K0701) and were subjected to Sanger sequencing (Macrogen,USA) using the sequencing primer 5′-GCAAACCCAAACTCCGAAGAC*-*3′. The degree of AEI for *CCL2* was expressed as a ratio of reference allele (REF; major) to alternative allele (ALT; minor). PeakPicker,v.2.0 was used with default settings to quantify the relative amount of the two alleles from the chromatogram after peak intensity normalization (Ge et al., 2005).

### Bioinformatic analyses

The ex vivo binding of RBPs to region flanking rs13900 and *CCL2* 3′ UTR was examined using Atlas of UTR Regulatory Activity (AURA) (Dassi et al., 2014). The resources included in POSTAR3 were used to interrogate the RNA binding motifs that overlap the rs13900 (Zhao et al., 2022). RBP-var2 was used to explore the potential functional consequences (Mao et al., 2016), and the ViennaRNA package (Lorenz et al., 2011) was used to predict changes in *CCL2* secondary structure due to rs13900.

### RNA electrophoretic mobility shift assay (REMSA)

Oligoribonucleotides spanning the rs13900 C or rs13900 T alleles (IDT) labeled with infrared dyes were used to determine the differential binding of HuR. The sequences of the oligoribonucleotide with rs13900 C allele is rCrUrUrUrCrCrCrCrArGrArCrArCrCr**<u>C</u>**rUrGrUrUrUrUrArUrUrU and oligoribonucleotide with rs13900 T allele is rCrUrUrUrCrCrCrCrArGrArCrArCrCr**<u>U</u>**rUrGrUrUrUrUrArUrUrU. The oligoribonucleotides were incubated with either 10 µg HuR overexpressing HeLa cell whole cell extracts or 25-200 µM ELAVL1 human recombinant protein (Origene, Cat No: TP301562) in a 20 µL reaction mixture with 20 units of RNasin and 1X RNA binding buffer containing 20 mM HEPES (pH 7.6), 3 mM MgCl2, 40 mM KCl, 5% glycerol and 2 mM dithiothreitol, 20 µg yeast tRNA. The reaction was incubated on ice for 10 min and super-shift assay was performed by adding 2 µL of anti-HuR antibody (3A2, mouse monoclonal IgG antibody, Santa Cruz Biotechnology, Cat No: sc-5261). After antibody addition, the complexes were incubated on ice for 15 min and were resolved by electrophoresis under non-denaturing conditions in 5% polyacrylamide gels with 0.5 × TBE running buffer. Gels were then analyzed using Li-Cor Odyssey CLX.

### RNA-binding protein immunoprecipitation

RNA-binding protein immunoprecipitation (RIP) was carried out using an immunoprecipitation kit (RIP-assay kit, MBL) following the manufacturer’s instructions. Briefly 2x10^6^ cells were lysed and the extract was precleared and incubated with 15 μg of antibody against HuR or IgG for 3 h. RNA on the beads was purified and converted to cDNA using random hexamer. cDNA was further amplified and the PCR product sequenced using the following primer: GCAAACCCAAACTCCGAAGAC.

### Cell culture and transfection

HEK-293 cell line was maintained in EMEM (ATCC) containing 10% heat-inactivated FBS, penicillin (100 U/mL) and streptomycin (100mg, mL) (GIBCO). Cells were cultured at 37°C in humidified air containing 5% CO2. For HuR overexpression and silencing studies, cells were seeded in six-well plates in serum-free EMEM at a density of ∼ 250,000 cells per well. The cells were transfected when they reached ∼ 70 to 80% confluency, using lipofectamine 3000 (ThermoFisher Scientific, Cat No: L3000008) according to the manufacturer’s recommended protocol. Briefly, lipofectamine 3000 was diluted in OptiMEM and allowed to complex with either 0.5 µg of pCMV6-HuR or pCMV-Entry plasmid or 50 µM of HuR siRNA or control siRNA for 15 min at room temperature. The plasmid/siRNA-lipofectamine 3000 mixture was then added to the cells in a final volume of 2 mL of complete medium. After incubation for 72 h, cells were harvested and used for further processing.

### Western blotting

Cell lysates were prepared in chilled RIPA buffer (25 mM Tris-HCL pH 7.6, 150 mM NaCl, 1% NP-40, 1% sodium deoxycholate, 0.1% SDS; Thermo Scientific, Rockford, IL) containing protease inhibitor (complete Mini, EDTA-free Protease inhibitor cocktail tablets, Roche Diagnostics, Cat No: 11836170001). Cells were lysed on ice for 10-15 min followed by centrifugation at 12,000 × g for 15 min, and the cleared supernatant was collected and stored at −20°C. Cell lysates with equal amounts of total protein (15-20 μg) were loaded and separated on a SDS polyacrylamide gel (10–15%) and were electrophoretically transferred onto nitrocellulose membranes (Thermo Scientific). The membranes were blocked with non-fat dry milk in TBST (50 mM Tris-HCl, pH 7.4, 150 mM NaCl, 0.2% Tween 20) and were incubated overnight with anti-HuR primary antibodies (diluted 1:1000). Following incubation, the membranes were washed and then further incubated with anti-mouse ß-actin for one h at room temperature, followed by HRP labeled goat anti-mouse or goat anti-rabbit antibodies (1:1000; Santa Cruz). Protein bands were detected using a chemiluminescence (ECL kit) method (Pierce) and visualized on X-ray film (Kodak).

### Reporter assays

Site-specific mutagenesis was done on *CCL2* LightSwitch 3′ UTR reporter (Active Motif) to generate constructs containing rs13900 C or rs13900 T alleles. The nucleotide sequence of the complete constructs was verified by Sanger sequencing (Macrogen, USA). 0.5 µg of these constructs were nucleofected into HEK293 cells using 4D Nucleofector (Lonza) and luciferase activity was measured 24 h post transfection using Spectramax plate reader (Molecular Devices). The efficiency of the nucleofection was verified by confocal microscopy, and it was greater than 90%. Given the high efficiency of the nucleofection and to avoid cross-talk with co-transfected control plasmids (Farr and Roman, 1992), luciferase data across samples was normalized using the protein content of the lysates using BCA protein assay. mRNA translatability was assessed using methods as described previously (Zhang et al., 2017).

### Polysome profiling

Human monocytes were isolated from fresh blood as described earlier (Gavrilin et al., 2009) with slight modification. Briefly, peripheral blood mononuclear cells were isolated by density gradient centrifugation using Histopaque, followed by immunomagnetic negative selection using EasySep Human Monocyte isolation kit. A high purity level for CD14^+^ cells was consistently achieved (≥90%) through this procedure, as confirmed by flowcytometry. The purified monocytes were immediately used for macrophage differentiation by treating them with 50 ng/mL M-CSF (PeproTech) for 72 h and flow cytometric measurement of surface markers CD64+, CD206+, CD44 was used to confirm the differentiation (Fig. S6). Cells were treated with LPS or PBS for 3 h to induce *CCL2* expression. Cells were treated with 0.1 mM cycloheximide, a translation inhibitor, for 3 min, harvested and washed with ice-cold PBS containing 0.1 mM cycloheximide, and pelleted by centrifugation for 5 min at 500 x g. The pellets were either stored at −80°C or immediately used for cytoplasmic lysate preparation. The cell pellet was resuspended in 500 µL lysis buffer containing 150 mM NaCl, 50 mM Tris-HCl pH 7.5, 10 mM KCl, 10 mM MgCl_2_, 0.2% NP-40, 2 mM dithiothreitol, 2 mM sodium orthovanadate, 1 mM phenylmethylsulfonyl fluoride, and 80 units/mL RNaseOUT. After 10 min incubation on ice with occasional inverting every two minutes, samples were centrifuged at 12,000 x g for 10 min at 4°C to pellet the nuclei and the post-nuclear supernatants. The optical density at 254 nm was measured, and volumes corresponding to the same OD254 were used. Post-nuclear supernatants were laid on top of a 10-50% sucrose gradient. The gradient was centrifuged for 90 min at 200,000 x g. After the ultracentrifugation, ten fractions were collected from top to bottom. RNA was extracted from each fraction and RT-qPCR was performed using *CCL2* mRNA specific primers. For analysis, fraction one was considered as cytoplasmic lysate, fractions 2-4 were considered monosomal fraction, and fractions 6-10 as polysomal fraction. The percent (%) distribution for the *CCL2* mRNAs across the gradients was calculated using the differences in the cycle threshold (ΔCt) values using the following formula (Nayak et al., 2013): % of mRNA A in each fraction = 2^ΔCT^ ^fraction^ ^X^ x 100/Sum, where ΔCT fraction X = CT (fraction 1) − CT (fraction X)

### Lentiviral transduction of primary macrophages

Cells were transfected by slightly modifying the method described by Plaisance-Bonstaff et.al 2019 (Plaisance-Bonstaff et al., 2019). Briefly, monocytes were purified from PBMCs obtained from homozygous donors for rs13900 C or rs13900T by negative selection. Upon purification cells were resuspended in 24 well plates at a seeding density of 0.5 x10^6^ cells per well and were further cultured in the medium supplemented with 50 ng/mL M-CSF (Figure. S7). After 24 h, ready to use GFP-tagged pCMV6-HuR or CMV-null lentiviral particles (Amsbio, Cambridge, M.A) were transduced into 0.5 x10^6^ cells in presence of polybrene (60 µg/mL) at a MOI of 1. The cells were processed for HuR and *CCL2* expression 72 h after transduction after stimulation with LPS for 3 h.

### Statistical Analyses

All values in figures are presented as the mean ± SEM. Results were analyzed with ANOVA and posthoc contrasts with Fisher (Least Significant Difference (LSD) test or Student’s t test. Statistical analyses were carried out in the Sigma Plot 12.0 software.

## Acknowledgments

We thank the participants of the study for their time and support.

## Description of the supplemental data

The Supplemental data includes Supplementary Tables 1-4 and Figures S1-S7.

## Web resources

AURA, http://aura.science.unitn.it/ POSTAR3, http://postar.ncrnalab.org

RBP-var2, http://159.226.67.237/sun/RBP-Var/index.php/Rbp/index ViennaRNA package, https://www.tbi.univie.ac.at/RNA/

---

**Figure S1.**
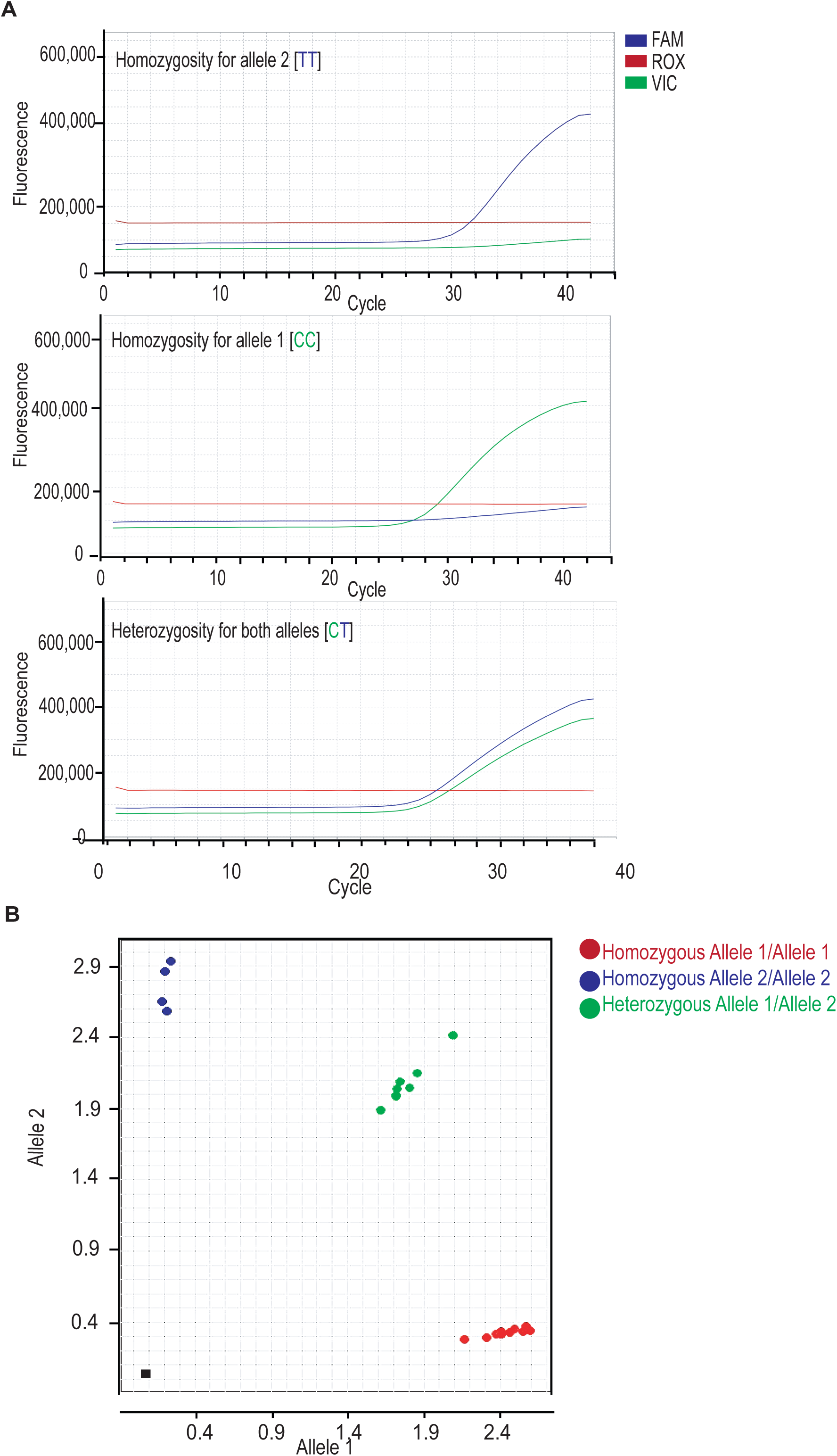
SNP genotyping for rs13900 using TaqMan technology. **(A)** Real-time multicomponent amplification curves for the TaqMan’s rs13900 assay. Fluorescence signal detected for individuals with rs13900T, rs13900C and rs13900CT genotypes. The blue curve indicates amplification of T allele (alternate/mutant allele) while the green curve represents C allele (wild type). **(B)** Allelic discrimination plot generated on Quant Studio 12K Flex Real-Time PCR system showing higher signal and better cluster separation for the rs13900 assay. Allele C is plotted on X-axis and the T allele is plotted on Y-axis.

**Figure S2.**
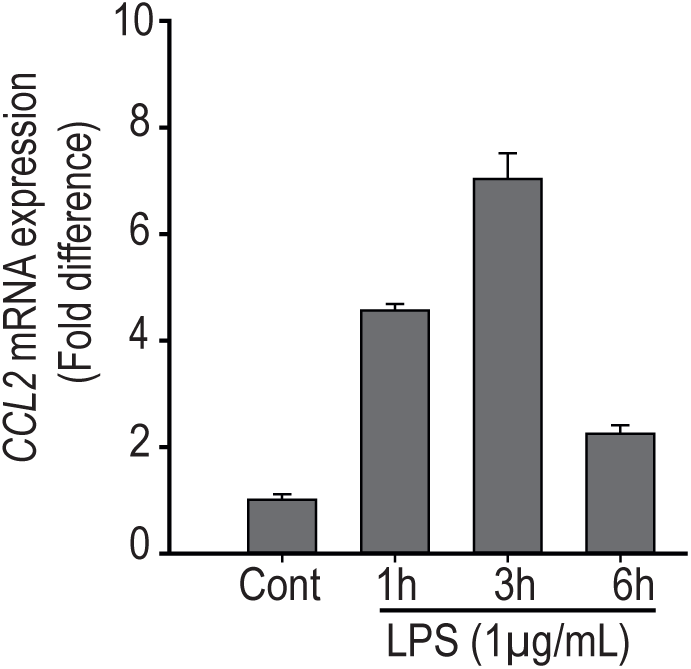
Time course of *CCL2* mRNA expression in PBMCs. Peripheral blood mononuclear cells (PBMC) were treated with 1 µg lipopolysaccharide (LPS) for 1, 3 or 6 hours. *CCL2* mRNA expression was measured by quantitative real-time PCR and normalized to 18S rRNA. Data are presented as mean ± SD of triplicate samples.

**Figure S3.**
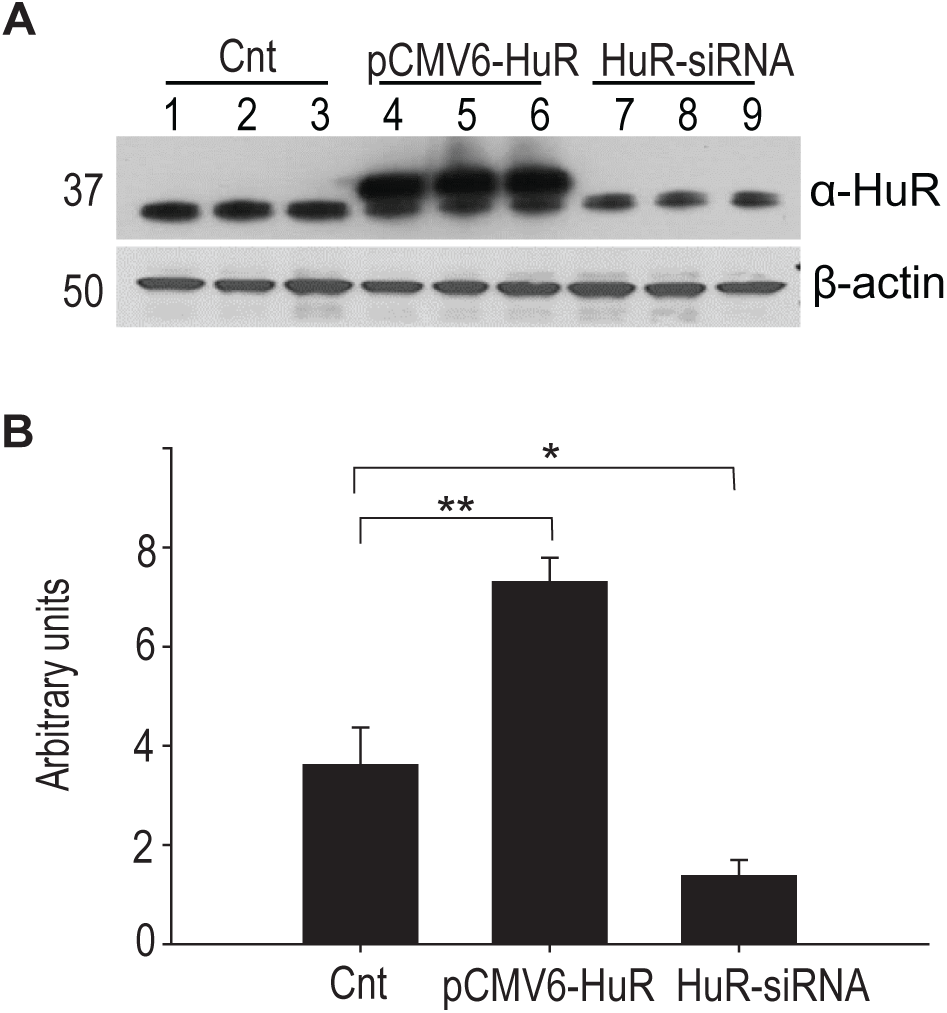
Validation of HuR overexpression and silencing by Western blotting. **(A)** A representative western blot showing HuR protein levels in HEK293 cells transfected with either pCMV6-HuR plasmid or HuR-targeting siRNA (HuR-siRNA). **(B)** Densitometric analysis of Western blot bands. The histogram shows relative intensity of HuR bands in each sample, normalized to β-actin. Error bar represents standard error mean from three independent experiments. *p<0.025, ** p<0.05.

**Figure S4.**
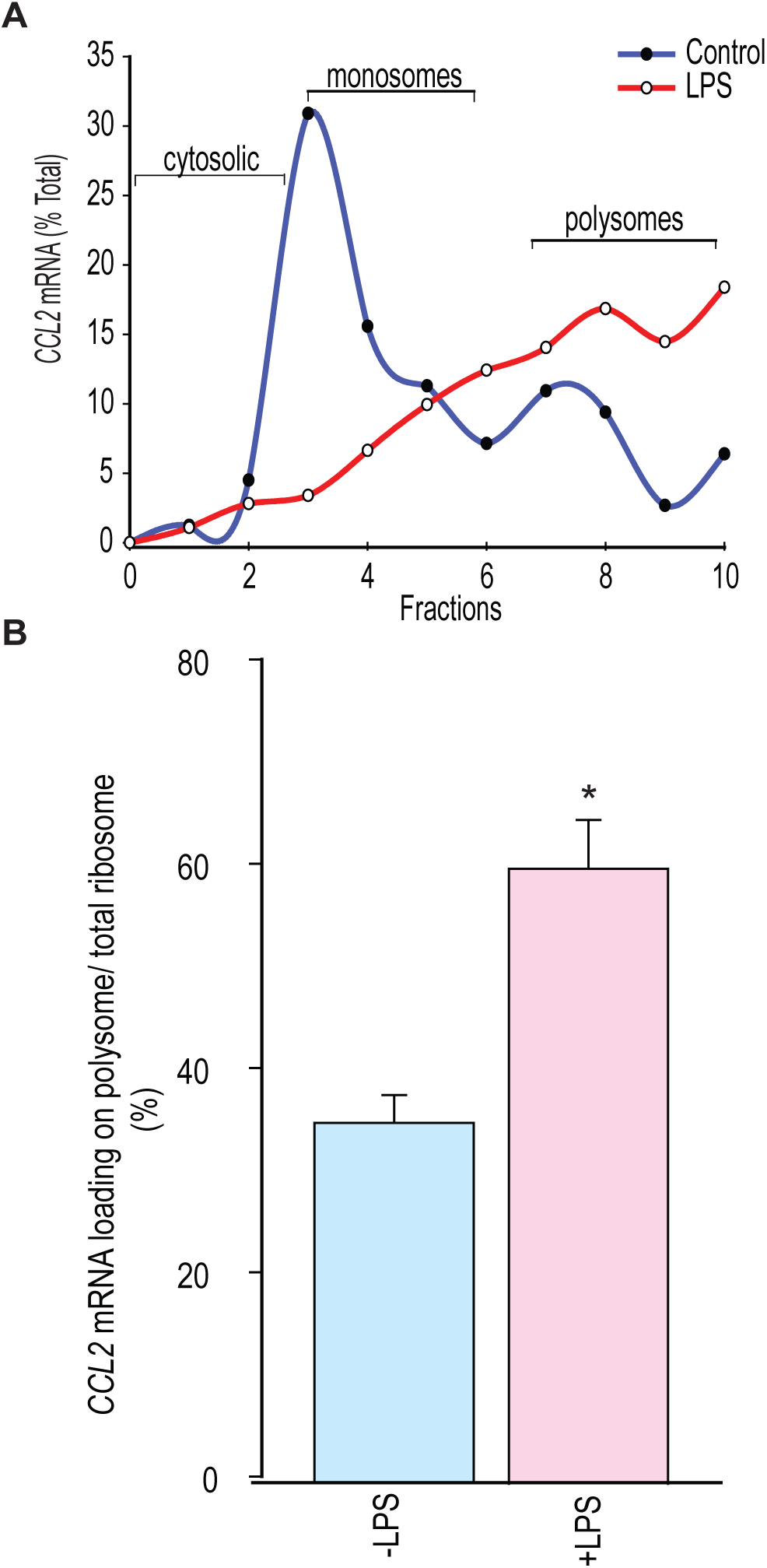
Polysomal association of *CCL2* mRNA before and after LPS stimulation of macrophages. **(A)** Polysome profile obtained by sucrose gradient centrifugation from macrophages before and after stimulation with LPS (1 µg/mL) for 3 h. The polysome profile shows a shift from monosome to heavier polysome upon LPS stimulation, indicating active translation. **(B)** The percentage of *CCL2* mRNA loading on polysome fraction was calculated using ΔCT method (mean ± SEM, n=4). *p<0.025.

**Figure S5.**
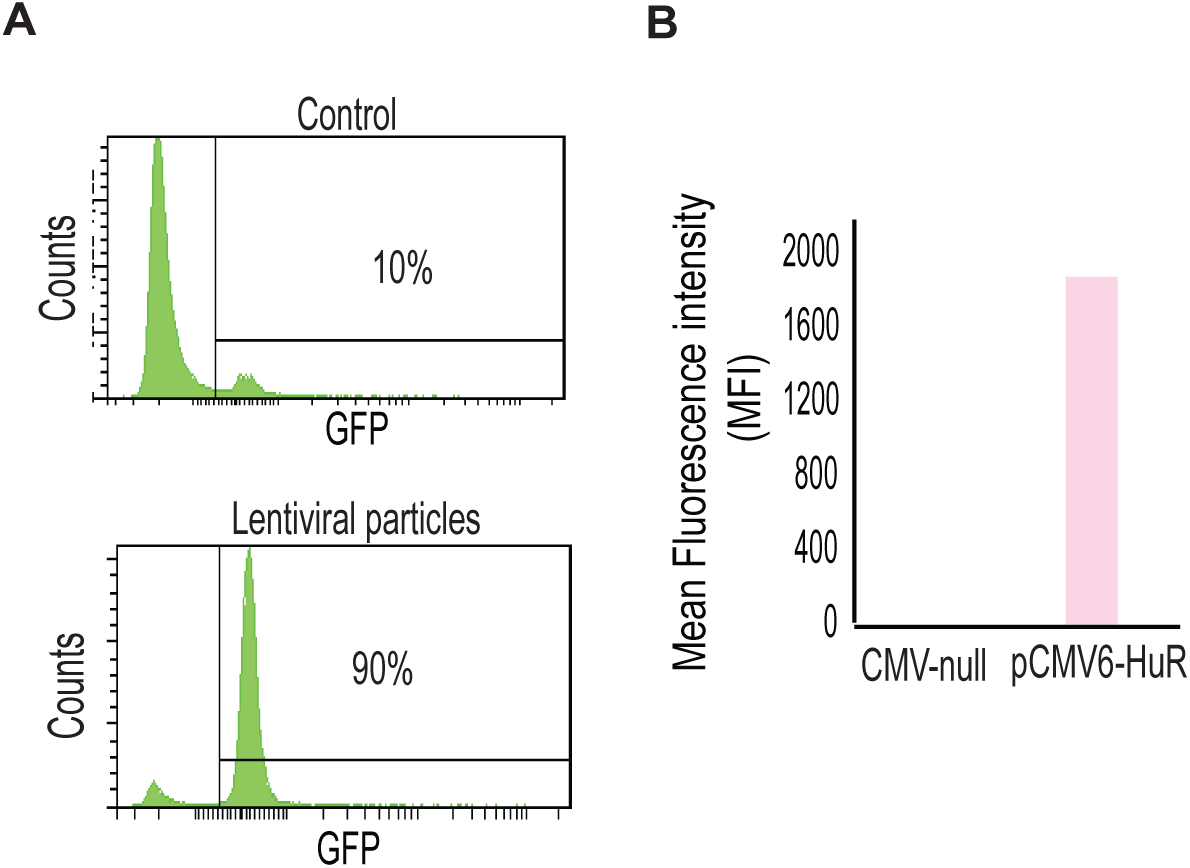
Transduction efficiency of pCMV6-HuR. Flow-cytometric analysis of transduction efficiency (Panel A) and the intensity of GFP in transduced cells (Panel B).

**Figure S6.**
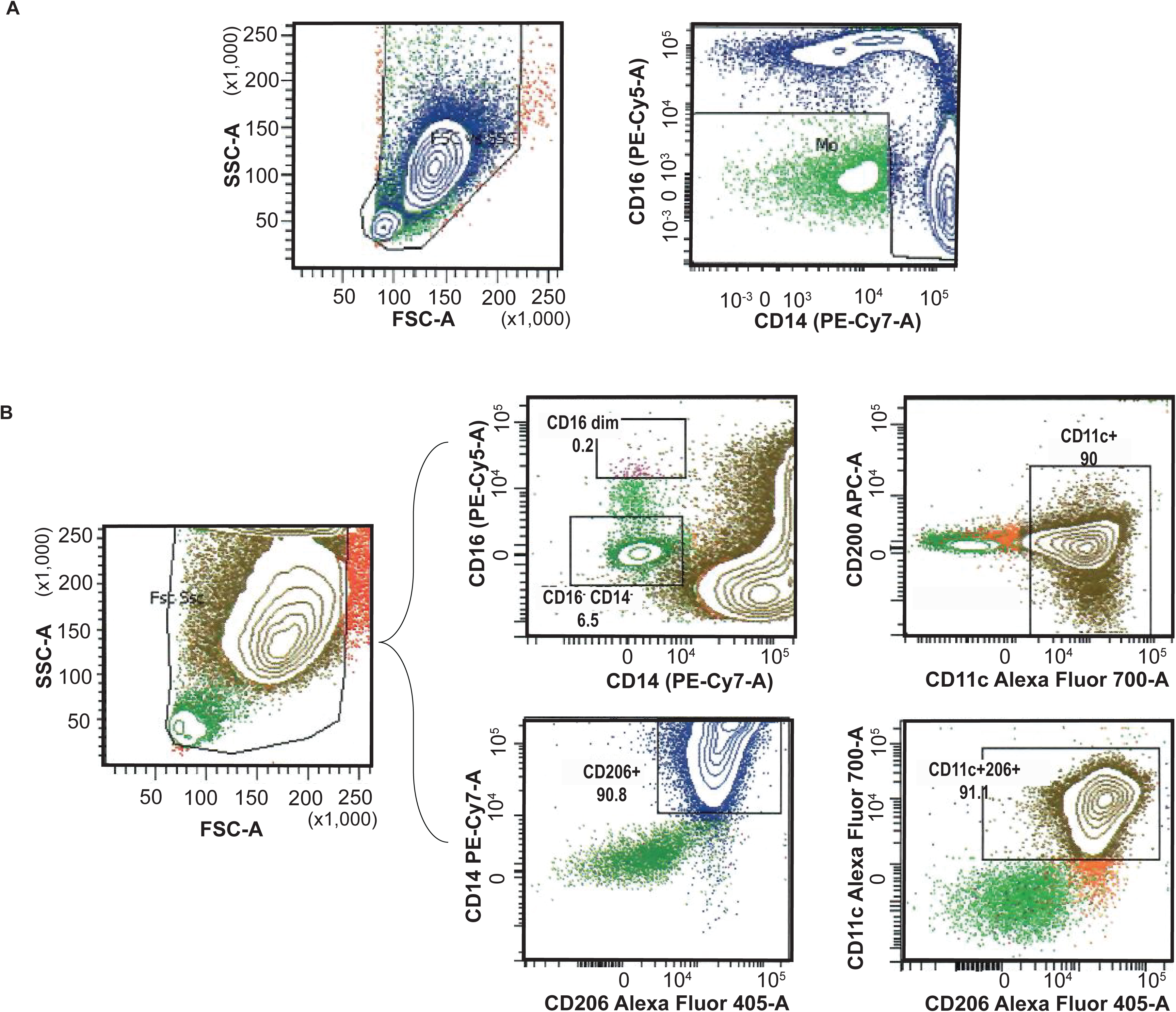
Flow cytometry analysis of cell surface markers to assess the purity of monocytes isolated from PBMC and in vitro differentiation to macrophages. **(A)** A representative image depicting the purity of monocytes isolated from PBMCs by immunomagnetic negative selection. A high monocyte content (>90%) was achieved in the enriched fraction. **(B)** Multicolor flow cytometry to determine macrophage differentiation. The macrophage population was identified by forward and side scatter and immunostaining. Surface expressions of CD11c, CD206, CD200, CD16 and CD14 were measured.

**Figure S7.**
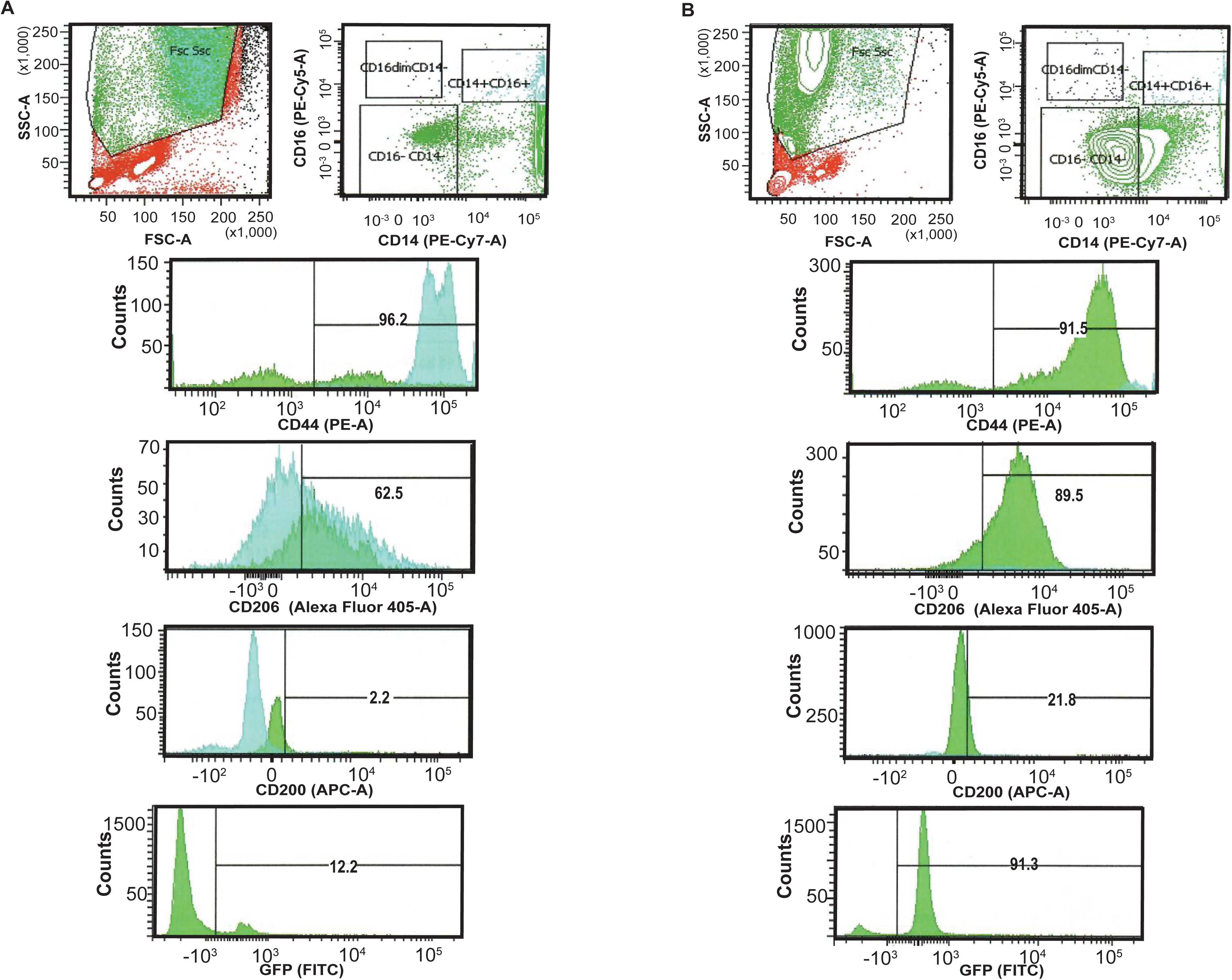
Lentiviral transduction of differentiated macrophages. The differentiation of monocytes into macrophages and their transduction was confirmed by detection of GFP and the cell surface markers CD44, CD200, CD206 by flow cytometry. Cells were transduced with either GFP-tagged pCMV-null **(A)** or GFP-tagged pCMV-HuR particles at MOI of 1 **(B)**. Each bracketed region in the histogram corresponds to the % of positive cells for respective markers noted on the x-axis of the histogram.

**Supplementary Table 1.** Allele carriages and allele frequencies of rs13900 in healthy volunteers, MAF, minor allele frequency.

|  | **rs13900**<br>**(n=42)** |
| --- | --- |
| Reference allele [CC] | 18 (42.8%) |
| Heterozygous [CT] | 16 (38.09%) |
| Alternative allele [TT] | 8 (19.04%) |
| MAF | 0.38 |
| Heterozygosity | 0.47 |
| No of alleles | 84 |
| Hardy Weinberg *P* | 0.459 |

**Supplementary Table 2.** Crosslinking immunoprecipitation (CLIP) analysis of HuR binding sites on the 3’untranslated region (3’UTR) of the *CCL2* gene. Summary of the SNPs located within HuR-binding regions is shown including SNP ID (rs#), genomic coordinates, strand, binding score, conservation scores (PhastCons, PhyloP), dataset accession numbers, alleles, and SNP position within peak region. Data was generated using POSTAR3 using the variation module.

| RBP | Tissue type | rs# | Position | Strand | CLIP-seq technology and peak calling method | Score | PhastCons score | PhyloP score | Data accession | Ref strand | Alt strand | SNP position |
| --- | --- | --- | --- | --- | --- | --- | --- | --- | --- | --- | --- | --- |
| ELAVL1 | HeLa | rs13900 | chr17:34256881-34256901 | + | PAR-CLIP,Piranha_0.01 | 11 | 0.024 | 0.227 | GSE29943,GSM741175 | C | T | chr17:34256891-34256892 |
| ELAVL1 | HeLa | rs181021073 | chr17:34256857-34256876 | + | PAR-CLIP,PARalyzer | 0.647 | 0.002 | 0.14 | GSE29943,GSM741173 | T | G | chr17:34256868-34256869 |
| ELAVL1 | HeLa | rs181021073 | chr17:34256857-34256876 | + | PAR-CLIP,PARalyzer | 0.653 | 0.002 | 0.14 | GSE29943,GSM741174 | T | G | chr17:34256868-34256869 |
| ELAVL1 | HeLa | NA | chr17:34257108-34257130 | + | PAR-CLIP,PARalyzer | 0.623 | 0.001 | -0.256 | GSE29943,GSM741173 | AT | A | chr17:34257109-34257110 |
| ELAVL1 | HeLa | NA | chr17:34257101-34257121 | + | PAR-CLIP,Piranha_0.01 | 9 | 0 | -0.39 | GSE29943,GSM741173 | AT | A | chr17:34257109-34257110 |
| ELAVL1 | HeLa | NA | chr17:34257101-34257121 | + | PAR-CLIP,Piranha_0.01 | 6 | 0 | -0.39 | GSE29943,GSM741175 | AT | A | chr17:34257109-34257110 |

**Supplementary Table 3.** Loading of rs13900 alleles to cytosolic, monosomal and polysomal fractions from macrophage extracts prepared from heterozygous donors. A T:C ratio > 1 indicates increased levels of the T allele relative to C allele.

|  | gDNA | Cytosol | Monosome | Polysome |
| --- | --- | --- | --- | --- |
| D-1 | 0.975806 | 0.872946 | 0.873425 | 1.778429 |
| D-2 | 1.034091 | 1.557778 | 1.441919 | 1.562622 |

**Supplementary Table 4.**
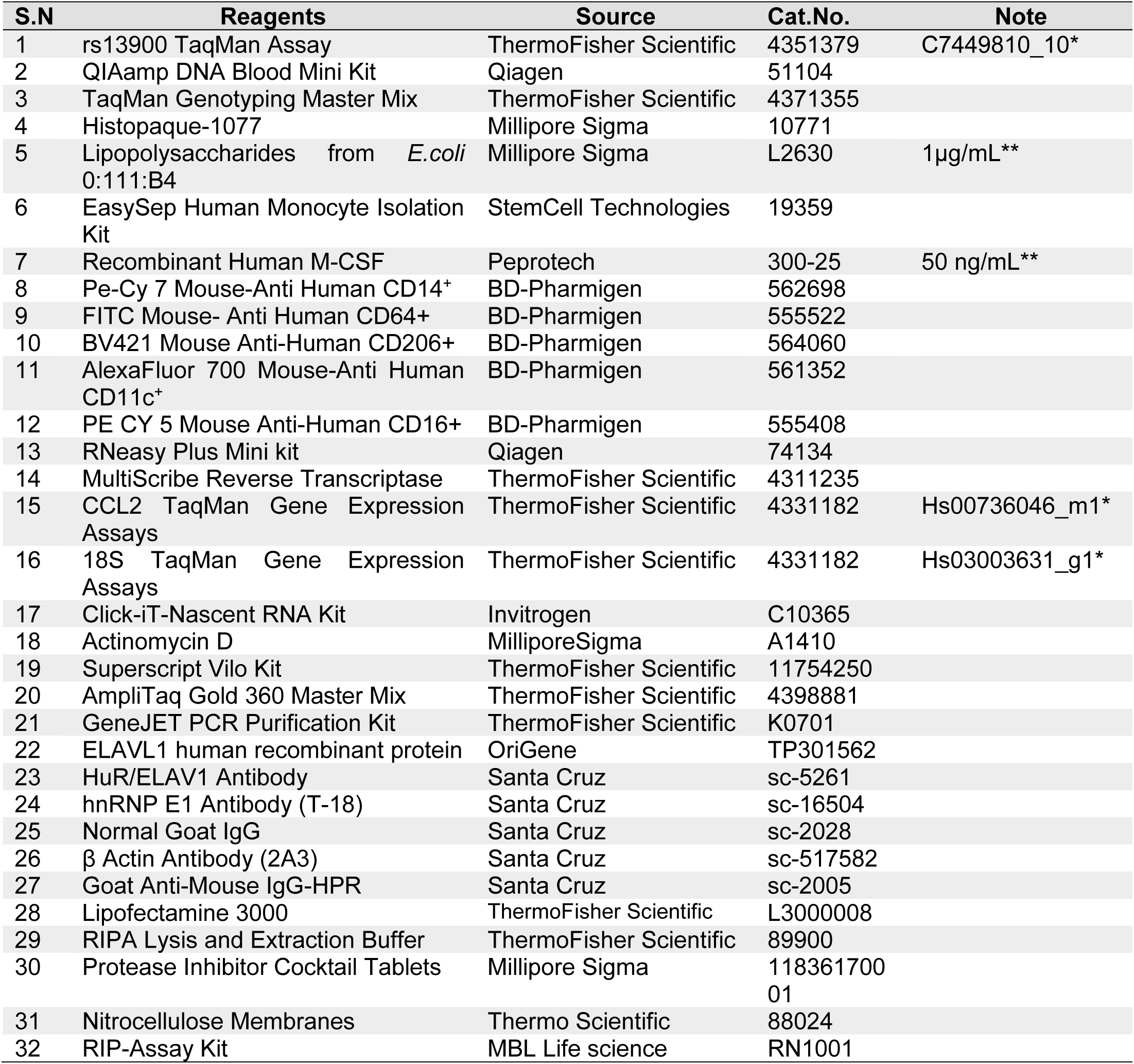
Reagents used in the study with source and catalog numbers.

## References

Akdeli, N., Riemann, K., Westphal, J., Hess, J., Siffert, W., Bachmann, H.S., 2014. A 3’UTR polymorphism modulates mRNA stability of the oncogene and drug target Polo-like Kinase 1. Mol Cancer 13, 87.

Alasoo, K., Rodrigues, J., Mukhopadhyay, S., Knights, A.J., Mann, A.L., Kundu, K., Consortium, H., Hale, C., Dougan, G., Gaffney, D.J., 2018. Shared genetic effects on chromatin and gene expression indicate a role for enhancer priming in immune response. Nat Genet 50, 424–431.

Alonso-Villaverde, C., Coll, B., Parra, S., Montero, M., Calvo, N., Tous, M., Joven, J., Masana, L., 2004. Atherosclerosis in patients infected with HIV is influenced by a mutant monocyte chemoattractant protein-1 allele. Circulation 110, 2204–2209.

Anderson, P., 2010. Post-transcriptional regulons coordinate the initiation and resolution of inflammation. Nat Rev Immunol 10, 24–35.

Bonello, G.B., Pham, M.H., Begum, K., Sigala, J., Sataranatarajan, K., Mummidi, S., 2011. An evolutionarily conserved TNF-alpha-responsive enhancer in the far upstream region of human CCL2 locus influences its gene expression. J Immunol 186, 7025–7038.

Buccitelli, C., Selbach, M., 2020. mRNAs, proteins and the emerging principles of gene expression control. Nat Rev Genet 21, 630–644.

Buniello, A., MacArthur, J.A.L., Cerezo, M., Harris, L.W., Hayhurst, J., Malangone, C., McMahon, A., Morales, J., Mountjoy, E., Sollis, E., Suveges, D., Vrousgou, O., Whetzel, P.L., Amode, R., Guillen, J.A., Riat, H.S., Trevanion, S.J., Hall, P., Junkins, H., Flicek, P., Burdett, T., Hindorff, L.A., Cunningham, F., Parkinson, H., 2019. The NHGRI-EBI GWAS Catalog of published genome-wide association studies, targeted arrays and summary statistics 2019. Nucleic Acids Res 47, D1005–D1012.

Cavestro, G.M., Zuppardo, R.A., Bertolini, S., Sereni, G., Frulloni, L., Okolicsanyi, S., Calzolari, C., Singh, S.K., Sianesi, M., Del Rio, P., Leandro, G., Franze, A., Di Mario, F., 2010. Connections between genetics and clinical data: Role of MCP-1, CFTR, and SPINK-1 in the setting of acute, acute recurrent, and chronic pancreatitis. Am J Gastroenterol 105, 199–206.

Chen, S., Deng, C., Hu, C., Li, J., Wen, X., Wu, Z., Li, Y., Zhang, F., Li, Y., 2016. Association of MCP-1-2518A/G polymorphism with susceptibility to autoimmune diseases: a meta-analysis. Clin Rheumatol 35, 1169–1179.

Cho, M.L., Kim, J.Y., Ko, H.J., Kim, Y.H., Kim, W.U., Cho, C.S., Kim, H.Y., Hwang, S.Y., 2004. The MCP-1 promoter -2518 polymorphism in Behcet’s disease: correlation between allele types, MCP-1 production and clinical symptoms among Korean patients. Autoimmunity 37, 77–80.

Choi, H.J., Yang, H., Park, S.H., Moon, Y., 2009. HuR/ELAVL1 RNA binding protein modulates interleukin-8 induction by muco-active ribotoxin deoxynivalenol. Toxicol Appl Pharmacol 240, 46–54.

Chowdhury, P., Khan, S.A., 2017. Significance of CCL2, CCL5 and CCR2 polymorphisms for adverse prognosis of Japanese encephalitis from an endemic population of India. Sci Rep 7, 13716.

Darkoh, C., Comer, L., Zewdie, G., Harold, S., Snyder, N., Dupont, H.L., 2014. Chemotactic chemokines are important in the pathogenesis of irritable bowel syndrome. PLoS One 9, e93144.

Dassi, E., Re, A., Leo, S., Tebaldi, T., Pasini, L., Peroni, D., Quattrone, A., 2014. AURA 2: Empowering discovery of post-transcriptional networks. Translation (Austin) 2, e27738.

de Lange, K.M., Moutsianas, L., Lee, J.C., Lamb, C.A., Luo, Y., Kennedy, N.A., Jostins, L., Rice, D.L., Gutierrez-Achury, J., Ji, S.G., Heap, G., Nimmo, E.R., Edwards, C., Henderson, P., Mowat, C., Sanderson, J., Satsangi, J., Simmons, A., Wilson, D.C., Tremelling, M., Hart, A., Mathew, C.G., Newman, W.G., Parkes, M., Lees, C.W., Uhlig, H., Hawkey, C., Prescott, N.J., Ahmad, T., Mansfield, J.C., Anderson, C.A., Barrett, J.C., 2017. Genome-wide association study implicates immune activation of multiple integrin genes in inflammatory bowel disease. Nat Genet 49, 256–261.

Deshmane, S.L., Kremlev, S., Amini, S., Sawaya, B.E., 2009. Monocyte chemoattractant protein-1 (MCP-1): an overview. J Interferon Cytokine Res 29, 313–326.

Duan, J., Shi, J., Ge, X., Dolken, L., Moy, W., He, D., Shi, S., Sanders, A.R., Ross, J., Gejman, P.V., 2013. Genome-wide survey of interindividual differences of RNA stability in human lymphoblastoid cell lines. Sci Rep 3, 1318.

Fan, J., Ishmael, F.T., Fang, X., Myers, A., Cheadle, C., Huang, S.K., Atasoy, U., Gorospe, M., Stellato, C., 2011. Chemokine transcripts as targets of the RNA-binding protein HuR in human airway epithelium. J Immunol 186, 2482–2494.

Farh, K.K., Marson, A., Zhu, J., Kleinewietfeld, M., Housley, W.J., Beik, S., Shoresh, N., Whitton, H., Ryan, R.J., Shishkin, A.A., Hatan, M., Carrasco-Alfonso, M.J., Mayer, D., Luckey, C.J., Patsopoulos, N.A., De Jager, P.L., Kuchroo, V.K., Epstein, C.B., Daly, M.J., Hafler, D.A., Bernstein, B.E., 2015. Genetic and epigenetic fine mapping of causal autoimmune disease variants. Nature 518, 337–343.

Farr, A., Roman, A., 1992. A pitfall of using a second plasmid to determine transfection efficiency. Nucleic Acids Res 20, 920.

Fenoglio, C., Galimberti, D., Lovati, C., Guidi, I., Gatti, A., Fogliarino, S., Tiriticco, M., Mariani, C., Forloni, G., Pettenati, C., Baron, P., Conti, G., Bresolin, N., Scarpini, E., 2004. MCP-1 in Alzheimer’s disease patients: A-2518G polymorphism and serum levels. Neurobiol Aging 25, 1169–1173.

Flores-Villanueva, P.O., Ruiz-Morales, J.A., Song, C.H., Flores, L.M., Jo, E.K., Montano, M., Barnes, P.F., Selman, M., Granados, J., 2005. A functional promoter polymorphism in monocyte chemoattractant protein-1 is associated with increased susceptibility to pulmonary tuberculosis. J Exp Med 202, 1649–1658.

Franke, A., McGovern, D.P., Barrett, J.C., Wang, K., Radford-Smith, G.L., Ahmad, T., Lees, C.W., Balschun, T., Lee, J., Roberts, R., Anderson, C.A., Bis, J.C., Bumpstead, S., Ellinghaus, D., Festen, E.M., Georges, M., Green, T., Haritunians, T., Jostins, L., Latiano, A., Mathew, C.G., Montgomery, G.W., Prescott, N.J., Raychaudhuri, S., Rotter, J.I., Schumm, P., Sharma, Y., Simms, L.A., Taylor, K.D., Whiteman, D., Wijmenga, C., Baldassano, R.N., Barclay, M., Bayless, T.M., Brand, S., Buning, C., Cohen, A., Colombel, J.F., Cottone, M., Stronati, L., Denson, T., De Vos, M., D’Inca, R., Dubinsky, M., Edwards, C., Florin, T., Franchimont, D., Gearry, R., Glas, J., Van Gossum, A., Guthery, S.L., Halfvarson, J., Verspaget, H.W., Hugot, J.P., Karban, A., Laukens, D., Lawrance, I., Lemann, M., Levine, A., Libioulle, C., Louis, E., Mowat, C., Newman, W., Panes, J., Phillips, A., Proctor, D.D., Regueiro, M., Russell, R., Rutgeerts, P., Sanderson, J., Sans, M., Seibold, F., Steinhart, A.H., Stokkers, P.C., Torkvist, L., Kullak-Ublick, G., Wilson, D., Walters, T., Targan, S.R., Brant, S.R., Rioux, J.D., D’Amato, M., Weersma, R.K., Kugathasan, S., Griffiths, A.M., Mansfield, J.C., Vermeire, S., Duerr, R.H., Silverberg, M.S., Satsangi, J., Schreiber, S., Cho, J.H., Annese, V., Hakonarson, H., Daly, M.J., Parkes, M., 2010. Genome-wide meta-analysis increases to 71 the number of confirmed Crohn’s disease susceptibility loci. Nat Genet 42, 1118–1125.

Ge, B., Gurd, S., Gaudin, T., Dore, C., Lepage, P., Harmsen, E., Hudson, T.J., Pastinen, T., 2005. Survey of allelic expression using EST mining. Genome Res 15, 1584–1591.

Glisovic, T., Bachorik, J.L., Yong, J., Dreyfuss, G., 2008. RNA-binding proteins and post-transcriptional gene regulation. FEBS Lett 582, 1977–1986.

Gonzalez, E., Rovin, B.H., Sen, L., Cooke, G., Dhanda, R., Mummidi, S., Kulkarni, H., Bamshad, M.J., Telles, V., Anderson, S.A., Walter, E.A., Stephan, K.T., Deucher, M., Mangano, A., Bologna, R., Ahuja, S.S., Dolan, M.J., Ahuja, S.K., 2002. HIV-1 infection and AIDS dementia are influenced by a mutant MCP-1 allele linked to increased monocyte infiltration of tissues and MCP-1 levels. Proc Natl Acad Sci U S A 99, 13795–13800.

Gschwandtner, M., Derler, R., Midwood, K.S., 2019. More Than Just Attractive: How CCL2 Influences Myeloid Cell Behavior Beyond Chemotaxis. Front Immunol 10, 2759.

Hao, S., Baltimore, D., 2009. The stability of mRNA influences the temporal order of the induction of genes encoding inflammatory molecules. Nat Immunol 10, 281–288.

Heinz, S., Benner, C., Spann, N., Bertolino, E., Lin, Y.C., Laslo, P., Cheng, J.X., Murre, C., Singh, H., Glass, C.K., 2010. Simple combinations of lineage-determining transcription factors prime cis-regulatory elements required for macrophage and B cell identities. Mol Cell 38, 576–589.

Hollander, D., Naftelberg, S., Lev-Maor, G., Kornblihtt, A.R., Ast, G., 2016. How Are Short Exons Flanked by Long Introns Defined and Committed to Splicing? Trends Genet 32, 596–606.

Huang, Z., Shi, J., Gao, Y., Cui, C., Zhang, S., Li, J., Zhou, Y., Cui, Q., 2019. HMDD v3.0: a database for experimentally supported human microRNA-disease associations. Nucleic Acids Res 47, D1013–D1017.

Hubal, M.J., Devaney, J.M., Hoffman, E.P., Zambraski, E.J., Gordish-Dressman, H., Kearns, A.K., Larkin, J.S., Adham, K., Patel, R.R., Clarkson, P.M., 2010. CCL2 and CCR2 polymorphisms are associated with markers of exercise-induced skeletal muscle damage. J Appl Physiol (1985) 108, 1651–1658.

Intemann, C.D., Thye, T., Forster, B., Owusu-Dabo, E., Gyapong, J., Horstmann, R.D., Meyer, C.G., 2011. MCP1 haplotypes associated with protection from pulmonary tuberculosis. BMC Genet 12, 34.

Jao, C.Y., Salic, A., 2008. Exploring RNA transcription and turnover in vivo by using click chemistry. Proc Natl Acad Sci U S A 105, 15779–15784.

Johnson, A.D., Zhang, Y., Papp, A.C., Pinsonneault, J.K., Lim, J.E., Saffen, D., Dai, Z., Wang, D., Sadee, W., 2008. Polymorphisms affecting gene transcription and mRNA processing in pharmacogenetic candidate genes: detection through allelic expression imbalance in human target tissues. Pharmacogenet Genomics 18, 781–791.

Joven, J., Coll, B., Tous, M., Ferre, N., Alonso-Villaverde, C., Parra, S., Camps, J., 2006. The influence of HIV infection on the correlation between plasma concentrations of monocyte chemoattractant protein-1 and carotid atherosclerosis. Clin Chim Acta 368, 114–119.

Kasztelewicz, B., Czech-Kowalska, J., Lipka, B., Milewska-Bobula, B., Borszewska-Kornacka, M.K., Romanska, J., Dzierzanowska-Fangrat, K., 2017. Cytokine gene polymorphism associations with congenital cytomegalovirus infection and sensorineural hearing loss. Eur J Clin Microbiol Infect Dis 36, 1811–1818.

Kazan, H., Ray, D., Chan, E.T., Hughes, T.R., Morris, Q., 2010. RNAcontext: a new method for learning the sequence and structure binding preferences of RNA-binding proteins. PLoS Comput Biol 6, e1000832.

Kenny, E.E., Pe’er, I., Karban, A., Ozelius, L., Mitchell, A.A., Ng, S.M., Erazo, M., Ostrer, H., Abraham, C., Abreu, M.T., Atzmon, G., Barzilai, N., Brant, S.R., Bressman, S., Burns, E.R., Chowers, Y., Clark, L.N., Darvasi, A., Doheny, D., Duerr, R.H., Eliakim, R., Giladi, N., Gregersen, P.K., Hakonarson, H., Jones, M.R., Marder, K., McGovern, D.P., Mulle, J., Orr-Urtreger, A., Proctor, D.D., Pulver, A., Rotter, J.I., Silverberg, M.S., Ullman, T., Warren, S.T., Waterman, M., Zhang, W., Bergman, A., Mayer, L., Katz, S., Desnick, R.J., Cho, J.H., Peter, I., 2012. A genome-wide scan of Ashkenazi Jewish Crohn’s disease suggests novel susceptibility loci. PLoS Genet 8, e1002559.

Kim, S., Kim, S., Chang, H.R., Kim, D., Park, J., Son, N., Park, J., Yoon, M., Chae, G., Kim, Y.K., Kim, V.N., Kim, Y.K., Nam, J.W., Shin, C., Baek, D., 2021. The regulatory impact of RNA-binding proteins on microRNA targeting. Nat Commun 12, 5057.

Kwong, A., Boughton, A.P., Wang, M., VandeHaar, P., Boehnke, M., Abecasis, G., Kang, H.M., 2021. FIVEx: an interactive eQTL browser across public datasets. Bioinformatics. Lebedeva, S., Jens, M., Theil, K., Schwanhausser, B., Selbach, M., Landthaler, M., Rajewsky, N., 2011. Transcriptome-wide analysis of regulatory interactions of the RNA-binding protein HuR. Mol Cell 43, 340–352.

Letendre, S., Marquie-Beck, J., Singh, K.K., de Almeida, S., Zimmerman, J., Spector, S.A., Grant, I., Ellis, R., Group, H., 2004. The monocyte chemotactic protein-1 -2578G allele is associated with elevated MCP-1 concentrations in cerebrospinal fluid. J Neuroimmunol 157, 193–196.

Li, Y.I., van de Geijn, B., Raj, A., Knowles, D.A., Petti, A.A., Golan, D., Gilad, Y., Pritchard, J.K., 2016. RNA splicing is a primary link between genetic variation and disease. Science 352, 600–604.

Liu, J.Z., van Sommeren, S., Huang, H., Ng, S.C., Alberts, R., Takahashi, A., Ripke, S., Lee, J.C., Jostins, L., Shah, T., Abedian, S., Cheon, J.H., Cho, J., Dayani, N.E., Franke, L., Fuyuno, Y., Hart, A., Juyal, R.C., Juyal, G., Kim, W.H., Morris, A.P., Poustchi, H., Newman, W.G., Midha, V., Orchard, T.R., Vahedi, H., Sood, A., Sung, J.Y., Malekzadeh, R., Westra, H.J., Yamazaki, K., Yang, S.K., International Multiple Sclerosis Genetics, C., International, I.B.D.G.C., Barrett, J.C., Alizadeh, B.Z., Parkes, M., Bk, T., Daly, M.J., Kubo, M., Anderson, C.A., Weersma, R.K., 2015. Association analyses identify 38 susceptibility loci for inflammatory bowel disease and highlight shared genetic risk across populations. Nat Genet 47, 979–986.

Liu, Y., Beyer, A., Aebersold, R., 2016. On the Dependency of Cellular Protein Levels on mRNA Abundance. Cell 165, 535–550.

Lorenz, R., Bernhart, S.H., Honer Zu Siederdissen, C., Tafer, H., Flamm, C., Stadler, P.F., Hofacker, I.L., 2011. ViennaRNA Package 2.0. Algorithms Mol Biol 6, 26.

MacArthur, J., Bowler, E., Cerezo, M., Gil, L., Hall, P., Hastings, E., Junkins, H., McMahon, A., Milano, A., Morales, J., Pendlington, Z.M., Welter, D., Burdett, T., Hindorff, L., Flicek, P., Cunningham, F., Parkinson, H., 2017. The new NHGRI-EBI Catalog of published genome-wide association studies (GWAS Catalog). Nucleic Acids Res 45, D896–D901.

Maharshak, N., Hart, G., Ron, E., Zelman, E., Sagiv, A., Arber, N., Brazowski, E., Margalit, R., Elinav, E., Shachar, I., 2010. CCL2 (pM levels) as a therapeutic agent in Inflammatory Bowel Disease models in mice. Inflamm Bowel Dis 16, 1496–1504.

Manolio, T.A., 2010. Genomewide association studies and assessment of the risk of disease. N Engl J Med 363, 166–176.

Mao, F., Xiao, L., Li, X., Liang, J., Teng, H., Cai, W., Sun, Z.S., 2016. RBP-Var: a database of functional variants involved in regulation mediated by RNA-binding proteins. Nucleic Acids Res 44, D154–163.

Martin, J.C., Chang, C., Boschetti, G., Ungaro, R., Giri, M., Grout, J.A., Gettler, K., Chuang, L.S., Nayar, S., Greenstein, A.J., Dubinsky, M., Walker, L., Leader, A., Fine, J.S., Whitehurst, C.E., Mbow, M.L., Kugathasan, S., Denson, L.A., Hyams, J.S., Friedman, J.R., Desai, P.T., Ko, H.M., Laface, I., Akturk, G., Schadt, E.E., Salmon, H., Gnjatic, S., Rahman, A.H., Merad, M., Cho, J.H., Kenigsberg, E., 2019. Single-Cell Analysis of Crohn’s Disease Lesions Identifies a Pathogenic Cellular Module Associated with Resistance to Anti-TNF Therapy. Cell 178, 1493–1508 e1420.

Matsumiya, T., Ota, K., Imaizumi, T., Yoshida, H., Kimura, H., Satoh, K., 2010. Characterization of synergistic induction of CX3CL1/fractalkine by TNF-alpha and IFN-gamma in vascular endothelial cells: an essential role for TNF-alpha in post-transcriptional regulation of CX3CL1. J Immunol 184, 4205–4214.

Maurano, M.T., Humbert, R., Rynes, E., Thurman, R.E., Haugen, E., Wang, H., Reynolds, A.P., Sandstrom, R., Qu, H., Brody, J., Shafer, A., Neri, F., Lee, K., Kutyavin, T., Stehling-Sun, S., Johnson, A.K., Canfield, T.K., Giste, E., Diegel, M., Bates, D., Hansen, R.S., Neph, S., Sabo, P.J., Heimfeld, S., Raubitschek, A., Ziegler, S., Cotsapas, C., Sotoodehnia, N., Glass, I., Sunyaev, S.R., Kaul, R., Stamatoyannopoulos, J.A., 2012. Systematic localization of common disease-associated variation in regulatory DNA. Science 337, 1190–1195.

Mayr, C., 2019. What Are 3’ UTRs Doing? Cold Spring Harb Perspect Biol 11.

McDermott, D.H., Yang, Q., Kathiresan, S., Cupples, L.A., Massaro, J.M., Keaney, J.F., Jr., Larson, M.G., Vasan, R.S., Hirschhorn, J.N., O’Donnell, C.J., Murphy, P.M., Benjamin, E.J., 2005. CCL2 polymorphisms are associated with serum monocyte chemoattractant protein-1 levels and myocardial infarction in the Framingham Heart Study. Circulation 112, 1113–1120.

Melgarejo, E., Medina, M.A., Sanchez-Jimenez, F., Urdiales, J.L., 2009. Monocyte chemoattractant protein-1: a key mediator in inflammatory processes. Int J Biochem Cell Biol 41, 998–1001.

Morrison, A.R., Yarovinsky, T.O., Young, B.D., Moraes, F., Ross, T.D., Ceneri, N., Zhang, J., Zhuang, Z.W., Sinusas, A.J., Pardi, R., Schwartz, M.A., Simons, M., Bender, J.R., 2014. Chemokine-coupled beta2 integrin-induced macrophage Rac2-Myosin IIA interaction regulates VEGF-A mRNA stability and arteriogenesis. J Exp Med 211, 1957–1968.

Mummidi, S., Bonello, G.B., Ahuja, S.K., 2009. Confirmation of differential binding of Interferon Regulatory Factor-1 (IRF-1) to the functional and HIV disease-influencing -2578 A/G polymorphism in CCL2. Genes Immun 10, 197–198; author reply 199.

Nayak, B.K., Feliers, D., Sudarshan, S., Friedrichs, W.E., Day, R.T., New, D.D., Fitzgerald, J.P., Eid, A., Denapoli, T., Parekh, D.J., Gorin, Y., Block, K., 2013. Stabilization of HIF-2alpha through redox regulation of mTORC2 activation and initiation of mRNA translation. Oncogene 32, 3147–3155.

Nedelec, Y., Sanz, J., Baharian, G., Szpiech, Z.A., Pacis, A., Dumaine, A., Grenier, J.C., Freiman, A., Sams, A.J., Hebert, S., Page Sabourin, A., Luca, F., Blekhman, R., Hernandez, R.D., Pique-Regi, R., Tung, J., Yotova, V., Barreiro, L.B., 2016. Genetic Ancestry and Natural Selection Drive Population Differences in Immune Responses to Pathogens. Cell 167, 657–669 e621.

Pabis, M., Popowicz, G.M., Stehle, R., Fernandez-Ramos, D., Asami, S., Warner, L., Garcia-Maurino, S.M., Schlundt, A., Martinez-Chantar, M.L., Diaz-Moreno, I., Sattler, M., 2019. HuR biological function involves RRM3-mediated dimerization and RNA binding by all three RRMs. Nucleic Acids Res 47, 1011–1029.

Page, S.H., Wright, E.K., Jr., Gama, L., Clements, J.E., 2011. Regulation of CCL2 expression by an upstream TALE homeodomain protein-binding site that synergizes with the site created by the A-2578G SNP. PLoS One 6, e22052.

Pai, A.A., Cain, C.E., Mizrahi-Man, O., De Leon, S., Lewellen, N., Veyrieras, J.B., Degner, J.F., Gaffney, D.J., Pickrell, J.K., Stephens, M., Pritchard, J.K., Gilad, Y., 2012. The contribution of RNA decay quantitative trait loci to inter-individual variation in steady-state gene expression levels. PLoS Genet 8, e1003000.

Palmieri, O., Latiano, A., Salvatori, E., Valvano, M.R., Bossa, F., Latiano, T., Corritore, G., di Mauro, L., Andriulli, A., Annesec, V., 2010. The -A2518G polymorphism of monocyte chemoattractant protein-1 is associated with Crohn’s disease. Am J Gastroenterol 105, 1586–1594.

Paulsen, M.T., Veloso, A., Prasad, J., Bedi, K., Ljungman, E.A., Tsan, Y.C., Chang, C.W., Tarrier, B., Washburn, J.G., Lyons, R., Robinson, D.R., Kumar-Sinha, C., Wilson, T.E., Ljungman, M., 2013. Coordinated regulation of synthesis and stability of RNA during the acute TNF-induced proinflammatory response. Proc Natl Acad Sci U S A 110, 2240–2245.

Payne, J.L., Khalid, F., Wagner, A., 2018. RNA-mediated gene regulation is less evolvable than transcriptional regulation. Proc Natl Acad Sci U S A 115, E3481–E3490.

Pham, M.H., Bonello, G.B., Castiblanco, J., Le, T., Sigala, J., He, W., Mummidi, S., 2012. The rs1024611 regulatory region polymorphism is associated with CCL2 allelic expression imbalance. PLoS One 7, e49498.

Plaisance-Bonstaff, K., Faia, C., Wyczechowska, D., Jeansonne, D., Vittori, C., Peruzzi, F., 2019. Isolation, Transfection, and Culture of Primary Human Monocytes. J Vis Exp.

Pullmann, R., Jr., Abdelmohsen, K., Lal, A., Martindale, J.L., Ladner, R.D., Gorospe, M., 2006. Differential stability of thymidylate synthase 3’-untranslated region polymorphic variants regulated by AUF1. J Biol Chem 281, 23456–23463.

Quach, H., Rotival, M., Pothlichet, J., Loh, Y.E., Dannemann, M., Zidane, N., Laval, G., Patin, E., Harmant, C., Lopez, M., Deschamps, M., Naffakh, N., Duffy, D., Coen, A., Leroux-Roels, G., Clement, F., Boland, A., Deleuze, J.F., Kelso, J., Albert, M.L., Quintana-Murci, L., 2016. Genetic Adaptation and Neandertal Admixture Shaped the Immune System of Human Populations. Cell 167, 643–656 e617.

Rabani, M., Levin, J.Z., Fan, L., Adiconis, X., Raychowdhury, R., Garber, M., Gnirke, A., Nusbaum, C., Hacohen, N., Friedman, N., Amit, I., Regev, A., 2011. Metabolic labeling of RNA uncovers principles of RNA production and degradation dynamics in mammalian cells. Nat Biotechnol 29, 436–442.

Reyes, R., Alcalde, J., Izquierdo, J.M., 2009. Depletion of T-cell intracellular antigen proteins promotes cell proliferation. Genome Biol 10, R87.

Ripin, N., Boudet, J., Duszczyk, M.M., Hinniger, A., Faller, M., Krepl, M., Gadi, A., Schneider, R.J., Sponer, J., Meisner-Kober, N.C., Allain, F.H., 2019. Molecular basis for AU-rich element recognition and dimerization by the HuR C-terminal RRM. Proc Natl Acad Sci U S A 116, 2935–2944.

Ross, J., 1995. mRNA stability in mammalian cells. Microbiol Rev 59, 423–450.

Rovin, B.H., Lu, L., Saxena, R., 1999. A novel polymorphism in the MCP-1 gene regulatory region that influences MCP-1 expression. Biochem Biophys Res Commun 259, 344–348.

Sasaki, Y., Dehnad, A., Fish, S., Sato, A., Jiang, J., Tian, J., Schroder, K., Brandes, R., Torok, N.J., 2017. NOX4 Regulates CCR2 and CCL2 mRNA Stability in Alcoholic Liver Disease. Sci Rep 7, 46144.

Schott, J., Reitter, S., Philipp, J., Haneke, K., Schafer, H., Stoecklin, G., 2014. Translational regulation of specific mRNAs controls feedback inhibition and survival during macrophage activation. PLoS Genet 10, e1004368.

Schultz, C.W., Preet, R., Dhir, T., Dixon, D.A., Brody, J.R., 2020. Understanding and targeting the disease-related RNA binding protein human antigen R (HuR). Wiley Interdiscip Rev RNA 11, e1581.

Schwerk, J., Savan, R., 2015. Translating the Untranslated Region. J Immunol 195, 2963–2971.

Sharif, O., Bolshakov, V.N., Raines, S., Newham, P., Perkins, N.D., 2007. Transcriptional profiling of the LPS induced NF-kappaB response in macrophages. BMC Immunol 8, 1.

Shatoff, E., Bundschuh, R., 2020. Single nucleotide polymorphisms affect RNA-protein interactions at a distance through modulation of RNA secondary structures. PLoS Comput Biol 16, e1007852.

Steri, M., Idda, M.L., Whalen, M.B., Orru, V., 2018. Genetic variants in mRNA untranslated regions. Wiley Interdiscip Rev RNA 9, e1474.

Szalai, C., Kozma, G.T., Nagy, A., Bojszko, A., Krikovszky, D., Szabo, T., Falus, A., 2001. Polymorphism in the gene regulatory region of MCP-1 is associated with asthma susceptibility and severity. J Allergy Clin Immunol 108, 375–381.

Tam, V., Patel, N., Turcotte, M., Bosse, Y., Pare, G., Meyre, D., 2019. Benefits and limitations of genome-wide association studies. Nat Rev Genet 20, 467–484.

Tu, X., Chong, W.P., Zhai, Y., Zhang, H., Zhang, F., Wang, S., Liu, W., Wei, M., Siu, N.H., Yang, H., Yang, W., Cao, W., Lau, Y.L., He, F., Zhou, G., 2015. Functional polymorphisms of the CCL2 and MBL genes cumulatively increase susceptibility to severe acute respiratory syndrome coronavirus infection. J Infect 71, 101–109.

Tucci, M., Barnes, E.V., Sobel, E.S., Croker, B.P., Segal, M.S., Reeves, W.H., Richards, H.B., 2004. Strong association of a functional polymorphism in the monocyte chemoattractant protein 1 promoter gene with lupus nephritis. Arthritis Rheum 50, 1842–1849.

Uren, P.J., Burns, S.C., Ruan, J., Singh, K.K., Smith, A.D., Penalva, L.O., 2011. Genomic analyses of the RNA-binding protein Hu antigen R (HuR) identify a complex network of target genes and novel characteristics of its binding sites. J Biol Chem 286, 37063–37066.

Vilmundarson, R.O., Duong, A., Soheili, F., Chen, H.H., Stewart, A.F.R., 2021. IRF2BP2 3’UTR Polymorphism Increases Coronary Artery Calcification in Men. Front Cardiovasc Med 8, 687645.

Visscher, P.M., Wray, N.R., Zhang, Q., Sklar, P., McCarthy, M.I., Brown, M.A., Yang, J., 2017. 10 Years of GWAS Discovery: Biology, Function, and Translation. Am J Hum Genet 101, 5–22.

Wang, J., Pitarque, M., Ingelman-Sundberg, M., 2006. 3’-UTR polymorphism in the human CYP2A6 gene affects mRNA stability and enzyme expression. Biochem Biophys Res Commun 340, 491–497.

Wang, Z., Kayikci, M., Briese, M., Zarnack, K., Luscombe, N.M., Rot, G., Zupan, B., Curk, T., Ule, J., 2010. iCLIP predicts the dual splicing effects of TIA-RNA interactions. PLoS Biol 8, e1000530.

Wright, E.K., Jr., Page, S.H., Barber, S.A., Clements, J.E., 2008. Prep1/Pbx2 complexes regulate CCL2 expression through the -2578 guanine polymorphism. Genes Immun 9, 419–430.

Wu, X., Brewer, G., 2012. The regulation of mRNA stability in mammalian cells: 2.0. Gene 500, 10–21.

Xia, J., Bogardus, C., Prochazka, M., 1999. A type 2 diabetes-associated polymorphic ARE motif affecting expression of PPP1R3 is involved in RNA-protein interactions. Mol Genet Metab 68, 48–55.

Yoshimura, T., Yuhki, N., Moore, S.K., Appella, E., Lerman, M.I., Leonard, E.J., 1989. Human monocyte chemoattractant protein-1 (MCP-1). Full-length cDNA cloning, expression in mitogen-stimulated blood mononuclear leukocytes, and sequence similarity to mouse competence gene JE. FEBS Lett 244, 487–493.

Yoshinaga, M., Takeuchi, O., 2019. Post-transcriptional control of immune responses and its potential application. Clin Transl Immunology 8, e1063.

Zhai, Y., Zhong, Z., Chen, C.Y., Xia, Z., Song, L., Blackburn, M.R., Shyu, A.B., 2008. Coordinated changes in mRNA turnover, translation, and RNA processing bodies in bronchial epithelial cells following inflammatory stimulation. Mol Cell Biol 28, 7414–7426.

Zhang, F., Lupski, J.R., 2015. Non-coding genetic variants in human disease. Hum Mol Genet 24, R102–110.

Zhang, X., Chen, X., Liu, Q., Zhang, S., Hu, W., 2017. Translation repression via modulation of the cytoplasmic poly(A)-binding protein in the inflammatory response. Elife 6.

Zhao, W., Zhang, S., Zhu, Y., Xi, X., Bao, P., Ma, Z., Kapral, T.H., Chen, S., Zagrovic, B., Yang, Y.T., Lu, Z.J., 2022. POSTAR3: an updated platform for exploring post-transcriptional regulation coordinated by RNA-binding proteins. Nucleic Acids Res 50, D287–D294.

Zou, J., Hormozdiari, F., Jew, B., Castel, S.E., Lappalainen, T., Ernst, J., Sul, J.H., Eskin, E., 2019. Leveraging allelic imbalance to refine fine-mapping for eQTL studies. PLoS Genet 15, e1008481.

